# Niche-based selection and metabolic plasticity in the microbiome: Chronic press disturbances reconfigure the taxo-phenomic landscape of an industrialized riverine ecosystem

**DOI:** 10.64898/2026.05.21.726876

**Authors:** Neelam Devpura, Kunal R. Jain, Datta Madamwar

## Abstract

Riverine ecosystems particularly in industrialized environment are subjected to chronic press disturbances, resulting from the decadal release of synthetic organic compounds and other xenobiotics. While indigenous microbial communities are highly sensitive to such stressors, the resulting metabolic restructuring and functional reshaping of the microbiome, driven by these long-term anthropogenic pressures remains poorly characterized. In this study, a microbial ecology of Bhadar River flowing across the Jetpur Industrial Estate, (Jetpur) were studied. Using a cross-sectional comparative approach, soil/sediment samples were collected from the diverse polluted and non-polluted sites from the estate. The taxonomic profiling using 16S rRNA gene amplicon sequencing, taxo-phenomic shifts (through metaphenomics) was studied, while the functional potential of metabolic pathways was validated using high-resolution shot-gun metagenomic study. Due to prolong pollution, the samples were rich in sulphur (9809 to 12391 mg/L), where polluted samples were having elevated COD (2432 to 4150 mg/L) as well as BOD (1000 to 1420 mg/L) values, along with the presence of heavy metals (e.g., Fe, Mg). Results revealed a distinct taxonomic shift at both the bacterial and archaeal levels. In non-polluted sites Proteobacteria (33 to 57%) dominated along with Acidobacteria and Actinobacteria, with diverse genera like *Alcaligenes* and *Serratia.* Whereas, polluted sites exhibited marked increase in Bacteroidetes (13 to 29%), Firmicutes, and Synergistetes and genera like *Alkalitalea*, *Mesotoga* and *Desulfomicrobium*, reflecting anaerobic, fermentative, and sulfate-reducing phenotypes. The archaeal communities at polluted sites were dominated by Euryarchaeota (78 to 99%), specifically methanogenic genera of *Methanosaeta* and *Methanocalculus*, contrasting with the *Methanomassiliicoccus* dominance in non-polluted areas. The alpha-diversity was marginally higher in polluted sites (Shannon: 4.11 to 4.81 vs. 3.81 to 5.39 (non-polluted)), but beta-diversity underscored clear separation (94% variance explained by pollution). The shot-gun metagenomic analysis indicated a substantial enhancement in anaerobic metabolic capacities within the polluted microbiome, primarily in sulphur respiration (dissimilatory sulfate reduction), methanogenesis (elucidating biogenic pathways), along with nitrogen cycling (identifying key denitrification and ammonification genes). The polluted microbiome have developed the potential to metabolise/degrade complex aromatic compounds (pcaK for benzoate/protocatechuate transport) and heavy metal resistance. The strong positive co-occurrences among anaerobic phyla (Thermotogae, Synergistetes, Bacteroidetes) in polluted sites was established, indicating syntrophic interactions for xenobiotic metabolism. These findings provide a theoretical ecological model for perturbed industrial ecosystems, emphasizing the role of habitat selection in shaping microbial functional diversity and demonstrate the remarkable adaptation of autochthonous communities to persistent press disturbances.

## 1.0 Introduction

The input of an enormous amount of xenobiotic and synthetic organic compounds posed a significant potential disturbance to the indigenous microbiome in a pristine environment such as river ecosystem (Valentine et al., 2012). Since the microbial diversity of a riverine environment is huge, the *in situ* functional characterizations remain poorly understood (Saunders et al., 2016).

The industrially polluted environment (such as the Bhadar River flowing across the Jetpur Industrial Estate) receives repeated pulses of xenobiotic compounds throughout the year and for several decades (till date the anthropogenic activities continue). Because of such chronic release, the microbiome faces a press (long-term) disturbance (particularly concerning the time scale of their generation cycles and migration process) (Rodriguez-R et al., 2015). Chronic release and press-disturbance often found to result in microbial community shifts, driven by responses to individual traits and community actions. The changes occur presumably due to sensitivity towards toxic synthetic xenobiotic compounds (Rodriguez-R et al., 2015).

With few studies suggesting an insignificant effect of perturbation, others have shown the resistance and resilience of microbial processes post-disturbance, which may increase with increasing diversity (Shade et al., 2012a,b; Griffiths et al., 2013; Hartmann et al., 2014). Such diverging observations might be because of the prevailing of high functional redundancy in the soil microbiome. Studying the dynamics of microbial community diversity and functional characteristics in response to environmental changes is a major challenge (Kuang et al., 2016).

A dynamic pristine riverine ecosystem interconnecting with the terrestrial environment brings a high nutrient flux into the river, where obligate and facultative aerobic phenotypes preferentially prevail at the top strata of the water column. The amount of ‘carbon’ transported into the ocean by river, is merely a small fraction of the total carbon received by river from the terrestrial ecosystem. Because of indigenous microbial metabolism (through the carbon cycle), some of the ‘carbon’ is returned into the atmosphere in the form of CO_2_, while another fraction is stored in the river sediment (Wang et al., 2016; Breton-Deval et al., 2019). Additionally, some carbon may also incorporated into the trophic chain.

After the onset of pollution, rivers now carry and transport an unusually high amount of carbon and other nutrients (due to anthropogenic activities), along with the carbon from terrestrial habitats. Therefore, a perturbed river becomes a further complex and heterogeneous ecosystem, since it now contains a high amount of organic carbons, substituted and non-substituted aromatic compounds, hydrocarbons, halogenated compounds, organophosphates (in form of fertilizers and pesticides), heavy metals and other xenobiotic compounds. As a consequence, the biogeochemistry and nutrient flux of the polluted river is unbalanced, which is because the energy captured by the system is unusually higher than its requirement. Moreover, due to the persistent and chronic pollution, the microbial phenotype has been skewed towards anaerobic respiration.

The mechanism and principle of the assemblage of microbiome after perturbations (such as the input of xenobiotic compounds into the open environment) is a multi-factorial event and yet to be elucidated completely. However, such niche-based selection is likely to have a role in shaping the microbiome, with characteristic diversity and specific functions (Nannipieri et al., 2003). For the river sediment microbiome, the diversity-functioning relationship is not well understood. However, with recent development in high throughput molecular technologies, studies establishing the significance of microbial diversity in soil ecosystem functioning and stability are increasing (Ren et al., 2019a,b; Singh, 2016; Saleem et al., 2015).

With the above reference, we designed a study to understand the microbial ecology of the perturbed riverine ecosystem of Bhadar River (flowing across the Jetpur Industrial Estate, JIE). An attempt was made to elucidate the functional competence of the microbial community and its phylo-taxonomic relationship. The study was also aimed to predict the metaphenome prevailing at perturbed as well as the pristine ecosystem of JIE, Jetpur.

## 2.0 Materials and Methods

### 2.1 Study design, site description and sample collection

In the present study, a cross-sectional comparative analysis of taxo-phenomic and functional features of microbiomes of polluted (Bhadar River) (n = 5) and pristine/control (n = 5) sites of JIE was carried out. The spatial distribution of community assemblages and structure within the polluted as well as pristine sites were also analyzed. The interaction between microbial players and their metabolic contributions to the organizational pattern of ecosystem was predicted using imputed metagenomics and validated using shotgun whole metagenomic approach.

The study was carried out from contaminated samples collected near Bhadar River and pristine regions in the proximal areas under 500 m circumference in the JIE. The industrial units of the estate constantly (for the last 30 years) release liquid waste into the nearby canal flowing across the estate. Five distinct regions (points) were selected along the 1000 m transect of the canal, within JIE. The topography/geography of the canal was marked with distinct peculiarities: it has randomly distributed ditches (of few meters width and depth) and puddles, plain terrain, small top-surface elevations (of few inches) and falls, marshy, muskeg and swamp regions. The sampling points were selected such that soil and sediment samples from each distinct topography can be collected.

Wastewater regularly get accumulates in puddles and ditches over a time, pollutants accumulates and get concentrated in such locations. Conversely, in plain terrain and flat surfaces of the canal where wastewater flows constantly, the accumulation of xenobiotics is lower at such sites. However due to leaching effect, these compounds seep into the top surfaces of the soil beneath the flowing water.

Samples were collected a few inches away from the bank of the canal (visually equivalent to foreshore flats) in a stratified random manner. From each selected points three sub-samples were collected using spirit cleaned spade at a depth of ∼ 5 – 7 cm by digging 0.5 x 0.5 m square plots. The minimum distance between any two sampling points was at least 50 – 70 m. For the pristine site (sites were within the area of JIE, where visibly no pollution observed), a similar approach was applied and soil samples were collected from five distinct points. The collected soil and sediment samples were homogenized, plant debris, roots and other coarse particles were removed and metagenomic DNA was extracted within 24 h of sampling.

### 2.2 Metagenomic DNA extraction and sequencing

#### 2.2.1 DNA extraction

Metagenomic DNA from each sub-sample from polluted and pristine samples was independently extracted using the pre-standardized method of the lab (Desai and Madamwar, 2007) with few modifications. About twenty gram soil samples were mixed with extraction buffer (10% (w/v) sucrose, 1.5 M NaCl, 1% CTAB, 100 mM Tris-Cl (pH 8.0), 100 mM EDTA (pH 8.0) and 50mM sodium phosphate buffer), 0.5 g glass beads (1.5 – 2.0 mm) and incubated under shaking conditions (150 rpm) at 37°C for 20-30 min. This homogenous suspension was treated with Proteinase K (100 μg/ml) and lysozyme (10 mg/ml), incubated at 60°C for 30 min, followed by the addition of 1 g of PAC and further incubated for 30 min with intermittent mixing. The crude lysate was extracted with an equal volume of phenol:chloroform:isoamyl alcohol mixture (25:24:1) followed by a repeated (3-5 times) treatment with chloroform. Metagenomic DNA was precipitated with 0.8^th^ volume of isopropanol, washed with 70% ethanol, air-dried and resuspended in a minimum quantity of ultrapure water (Milli-Q grade). The extracted metagenomic DNA was purified using a silica-based gel extraction kit (Merck Biosciences, Bangalore, India) according to the manufacturer’s instructions. The quality and quantity of the extracted DNA were assessed using agarose gel electrophoresis and a nano-spectrophotometer, respectively.

#### 2.2.2 DNA sequencing

##### 2.2.2.1 16S rRNA gene amplicon sequencing

The extracted metagenomic DNA from polluted and pristine samples was used as a template for amplifying the V3-V4 region of the 16S rRNA gene of eubacteria. The obtained amplicons were sequenced using paired-end sequencing method on an Illumina MiSeq using 2 x 300 bp chemistry, as per the manufacturer’s instructions.

##### 2.2.2.2 Whole metagenome sequencing

For whole metagenome sequencing (shotgun approach), extracted metagenomic DNA from all sub-samples of polluted and pristine samples were independently mixed in equimolar ratios to form composite DNA representing the polluted and pristine ecosystems of JIE.

The composite DNA was sequenced by the paired-end sequencing method on Illumina Hiseq 4000 using 2 x 150 bp chemistry, as per the manufacturer’s instructions.

### 2.3 Post-sequencing analysis

The raw 16S rRNA amplicon sequenced reads were processed for adapter trimming using CutAdapt v.1.1.1 (Martin, 2011) followed by merging of paired-end reads using FLASH v.1.2.11 (Magoc and Salzberg, 2011). The low quality merged reads removed using SICKLE v.1.21 and chimeras were removed using ChimeraSlayer (Joshi and Fass, 2011). The obtained reads were clustered using cd-hit v.4.6 (identity >99%, coverage >80%) and de novo and closed referenced operational taxonomic units (OTU 97%) were picked using the QIIME 1.9.1 package (against Greengenes database (2013) (Caporaso et al., 2010). The reads were also annotated and analyzed using the RDP classifier v.2.11 and NCBI megablast by default parameters.

The metabolic functions of sequenced amplicons from both polluted and pristine samples were predicted using the Phylogenetic Investigation of Communities by Reconstruction of Unobserved States (PICRUSt) (Langille et al., 2013). To evaluate the accuracy of the predicted metagenome profiles, nearest sequenced taxon index (NSTI) scores were calculated. Gene content was assigned using the Kyoto Encyclopedia of Genes and Genomes (KEGG) Orthology (KO) copy numbers. Each predicted KO was subsequently fed to the KO database and binned into six main pathways (e.g., degradation of aromatic compounds, etc.) using the imputed metagenome.

From whole metagenome sequencing, raw sequenced reads were processed for adapter trimming, removal of low-quality reads, digital normalization to coverage (=20), followed by trimming of high-coverage reads at low-abundance k-mers and a second round of digital normalization to coverage (=5) using Khmer v.0.8.4. The high quality and digital normalized reads were assembled using a de novo approach using IDBA-UD v.1.1.1. (Peng et al., 2012). The assembled (meta)genome was used for gene prediction using Prokka v.1.10 (prokaryotic genome annotation tool) (Seemann, 2014). Genes from the assembled metagenome (scaffolds) were annotated using Uniprot (using search tool BLAST + blastp v.2.2.29+), RefSeq (search tool BLAST + blastp v.2.2.29+) and Pfam (search tool HMMER 3.1b1) databases. Moreover, the assembled metagenomic sequences of polluted and pristine samples were uploaded on MG-RAST server for automated annotation (Wilke et al., 2017).

### 2.4 Statistics

The microbial (bacterial) community structures were initially analyzed using alpha diversity indices (Shannon estimators, Chao1 and other indices). The similarity patterns amongst the samples were analyzed as principal coordinates analysis (PCoA) using two distance matrices, Bray-Curtis without considering microbial evolution and the unweighted UniFrac metric by accounting for the inferred evolutionary distance. The community similarity/dissimilarity was further analysed for (a) hierarchical clustering for bacterial community structure and composition, (b) nonparametric multivariate analysis of variance combined with analysis of similarities, (c) the overlapping in membership between communities and (d) core diversity was estimated using MetaComet (Wang et al., 2016) from default parameters. Calculations were based on both Bray-Curtis and Jaccard distance metrics, 1000 permutations and a false discovery rate (FDR) of 0.05 (Jordaan et al., 2019). PERMANOVA was applied to determine whether polluted and pristine bacterial communities harbour significantly different metagenomes (Jordaan et al., 2019).

## 3.0 Results

### 3.1 Physico-chemical characterization of sampling sites

The physico-chemical characteristics of each of the five sampling sites of polluted and pristine ecosystems were as described in the Table 1. The pH of the polluted samples at the time of sampling ranged from 7.27 to 8.10. The COD of polluted samples was expectedly higher ranging from 2432 to 4150 mg/L and BOD was 1000 to 1420 mg/L. The pristine habitat had low COD and BOD values, indicating the level of xenobiotic perturbations at polluted sites of Jetpur Industrial Estate. The sulphur concentration was ranging from 9,809 to 12,391 mg/L at polluted habitats. The total organic carbon and nitrogen concentrations also varied across the sampling sties (at polluted habitats), however was relatively less as compared to sulphur.

**Table 1:** Physico-chemical characteristics of polluted (JP-1 to JP-5) and pristine/non-polluted (JC-1 to JC-5) soil samples of Jetpur Industrial Estate, Jetpur.

| No. |  | JP-1 | JP-2 | JP-3 | JP-4 | JP-5 | JC-1 | JC-2 | JC-3 | JC-4 | JC-5 |
| --- | --- | --- | --- | --- | --- | --- | --- | --- | --- | --- | --- |
| **Physical Parameters** |  |  |  |  |  |  |  |  |  |  |  |
| 1 | pH | 7.40 | 7.27 | 7.42 | 7.70 | 8.10 | 7.81 | 8.34 | 8.54 | 8.00 | 8.12 |
| 2 | EC<br>(ms/ppt) | 8.895 | 7.589 | 7.956 | 6.985 | 7.015 | 2.874 | 0.925 | 0.589 | 1.541 | 0.945 |
| 3 | Salinity<br>(ppt) | 5.30 | 3.56 | 3.48 | 5.26 | 2.96 | 1.11 | 0.69 | 1.35 | 1.89 | 0.48 |
| 4 | CEC<br>(Meq/100g) | 345.12 | 359.47 | 386.21 | 289.58 | 169.85 | 111.63 | 98.63 | 128.00 | 198.68 | 124.23 |
| **Chemical Parameters** |  |  |  |  |  |  |  |  |  |  |  |
| 1 | COD<br>(mg/L) | 3050 | 3050 | 4150 | 3245 | 2432 | 222 | 156 | 170 | 170 | 163 |
| 2 | BOD<br>(mg/L) | 1400 | 1398 | 1420 | 1387 | 1000 | 101 | 90 | 93 | 102 | 92 |
| 3 | TOC<br>(%) | 11.569 | 13.569 | 18.017 | 09.874 | 23.012 | 1.256 | 0.984 | 0.910 | 0.894 | 0.648 |
| 4 | Na<br>(mg/kg) | 6547.12 | 5013.02 | 6984.65 | 3269.16 | 5687.24 | 157.01 | 346.89 | 192.46 | 500.38 | 482.21 |
| 5 | K<br>(mg/kg) | 3541 | 1142 | 954 | 8465 | 6873 | 117.14 | 93.01 | 145 | 102.34 | 95.39 |
| 6 | Ca<br>(mg/kg) | 6087.08 | 4684.5 | 7451.12 | 4035.74 | 7120.16 | 1046.01 | 3045.23 | 2541.32 | 1354.21 | 2647.10 |
| 7 | P<br>(mg/kg) | 10.21 | 12.06 | 8.77 | 9.56 | 11.38 | 2.54 | 1.24 | 0.73 | 0.53 | 1.87 |
| 8 | S<br>(mg/kg) | 11208 | 9980 | 11805 | 9809 | 12391 | 623 | 700 | 501 | 534 | 490 |
| 9 | Cl<br>(mg/kg) | 12045 | 13090 | 9771 | 19873 | 16988 | 1002 | 980 | 1121 | 1204 | 1416 |
| 10 | N<br>(%) | 15.4 | 31.6 | 21.2 | 17.2 | 26.03 | 7.40 | 6.54 | 4.24 | 2.8 | 12.47 |

As observed in Table 2, the various heavy metals concentrations were expectedly high from polluted samples as compared to pristine habitats, particularly iron and magnesium.

**Table 2:** Concentration of various heavy metals at polluted (JP-1 to JP-5) and pristine/non-polluted (JC-1 to JC-5) sites of Jetpur Industrial Estate, Jetpur.

| No. |  | JP-1 | JP-2 | JP-3 | JP-4 | JP-5 | JC-1 | JC-2 | JC-3 | JC-4 | JC-5 |
| --- | --- | --- | --- | --- | --- | --- | --- | --- | --- | --- | --- |
| 1 | Cr<br>(mg/kg) | 29.25 | 35.26 | 45.30 | 25.14 | 37.56 | 10.21 | 13.68 | 08.54 | 17.32 | BDL |
| 2 | Co<br>(mg/kg) | 8.12 | 3.57 | 13.08 | 7.87 | 15.00 | BDL | BDL | 10.25 | 6.84 | 7.06 |
| 3 | Fe<br>(mg/kg) | 5236.20 | 5142.56 | 4986.54 | 4963.11 | 5012.89 | 5623.23 | 5302.36 | 5912.08 | 5898.82 | 5176.24 |
| 4 | Cu<br>(mg/kg) | 48.01 | 36.20 | 155.12 | 43.05 | 51.45 | 25.10 | 22.02 | 45.25 | 36.87 | 39.23 |
| 5 | Mn<br>(mg/kg) | 558.11 | 503.62 | 389.48 | 432.17 | 500.078 | 723.12 | 723.82 | 635.47 | 584.64 | 610.68 |
| 6 | Zn<br>(mg/kg) | 488.12 | 402.35 | 394.25 | 189.05 | 387.99 | 62.87 | 50.36 | 93.07 | 55.83 | 90.30 |
| 7 | Cd<br>(mg/kg) | BDL | 2.37 | BDL | BDL | BDL | BDL | BDL | BDL | BDL | BDL |
| 8 | Ni<br>(mg/kg) | 17.60 | 23.07 | 14.17 | BDL | BDL | BDL | BDL | 3.70 | BDL | BDL |
| 9 | Mg<br>(mg/kg) | 8987.77 | 10532.5 | 9547.3 | 8215.3 | 10618.3 | 6852.6 | 5149.8 | 5610.5 | 6215.6 | 10245.8 |
| 10 | Hg<br>(mg/kg) | 12.58 | 14.23 | 12.36 | 13.48 | 12.09 | 05.42 | 06.65 | 07.77 | 10.35 | 10.74 |
| 11 | Pb<br>(mg/kg) | 50.16 | 38.10 | 46.27 | 40.44 | 60.08 | 47.90 | 26.33 | 20.04 | 36.00 | 37.92 |

BDL: Below detectable limit

### 3.2 Bacterial (microbial) diversity as a function of anthropogenic perturbation

To generate bacterial taxonomic profiles of sequenced reads of polluted and pristine samples OTUs with 97% similarity were clustered and annotated using Greengenes, RDP and NCBI databases. Each sequence was assigned to a taxonomic level using QIIME, RDP classifier and Blastn tools respectively. The classified sequences were analyzed for understanding the taxonomic structure and community diversity within and among the samples.

#### 3.2.1 Predicting microbial community composition using 16S rRNA amplicon sequencing

Microbial compositions in 10 soil samples across Jetpur polluted and pristine sites were generated by the MiSeq Illumina sequencing platform. Illumina sequencing generated the numbers of raw read in the range of 300,360 – 501,634 sequences. After removal of low-quality sequences and chimeric sequences, the numbers of analysed sequences per sample were in the range of 144,544 – 234,684 sequences. These pre-processed 16S rRNA sequences were then mapped and annotated based on NCBI, RDP databases as well as on Greengenes database on MG-RAST platform. The microbial community structure analysis was further divided into Archaeal and Eubacterial community composition.

#### 3.2.2. Bacterial Community Structure of Jetpur soil and soil/sediments

When analysing the community make up of Archaea and Eubacteria dominance of various phyla was differentially observed among all the amplicon sequences. All 10 soil samples showed a different relative abundance of several phyla, but upon grouping them all five of the pristine samples (JC-1 to JC-5), it showed abundance of Proteobacteria in the range of 33 % to 57 % making it the most abundant phylum. In contrast, the polluted sites (JP-1 to JP-5) showed a reduction in the abundance of Proteobacteria making about 21 % to 44 % (P = 0.035; FDR = 0.062). Members of the phyla Acidobacteria (5-16 %), Actinobacteria (5-16 %), Firmicutes (1.8 – 13.8 %), Chloroflexi (2- 10 %), Planctomycetes (2.6-6.4 %), Gemmatimonadetes (1.7 – 3.9 %) and Verrucomicrobia (1.7 – 2.5 %) were found to be most abundant at pristine samples after Proteobacteria. The relative abundance of these dominant phyla was making 90-95 % of the bacterial community at pristine samples.

While, bacterial community of polluted soil/sediment samples was showing majority of Bacteroidetes (∼13-29 %) (P = 0.003; FDR = 0.00056), Firmicutes (∼13-21 %) (P = 0.15; FDR = 0.2), Chloroflexi (∼2-23 %) (P = 0.24; FDR = 0.28), Synergistetes (∼0.9-12 %) (P = 0.00052; FDR = 0.0018), Thermotogae (∼0.8-14 %) (P = 0.0077; FDR = 0.18), Tenericutes (0.2-7%) (P = 0.008; FDR = 0.017), Verrucomicrobia (0.6-2.1 %) (P = 0.69; FDR = 0.71) and Spirochaetes (0.7– 3.6%) (P = 0.00052; FDR = 0.0018) after Proteobacteria. Members from these phyla were making 94-96% of the bacterial community at polluted soils. The relative abundance of these phyla was significantly different in each of the polluted sample, showing a micro-shift in the composition of these communities, which may indicate different types of pollutants across the 5 distinct sites (Figure 1).

**Figure 1:**
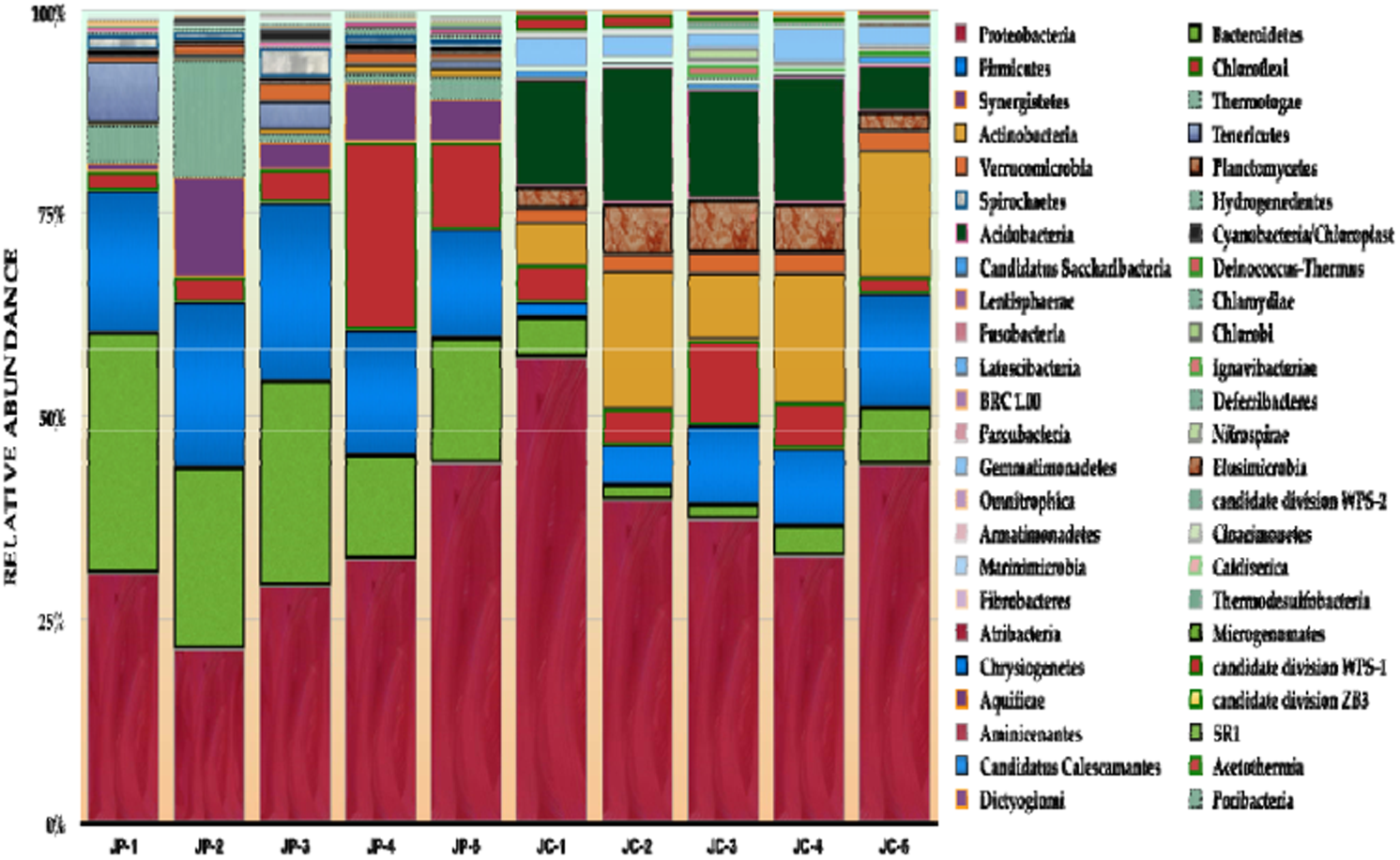
Differential abundance of bacterial community depicted at rank phylum among 10 sites of JIE mapped on RDP database. JP-1 to JP-5 represents polluted samples, JC-1 to JC-5 represents pristine/non-polluted samples.

When analysing the bacterial community structure at rank Class, significant shift was observed between the pristine and polluted community. At pristine site presence of various sub-divisions of Acidobacteria were observed making the community more diverse. Among most dominant classes were class Gammaproteobacteria (8-24 %) (P = 0.14; FDR = 0.17), Alphaproteobacteria (9.5-15 %) (P = 0.000037; FDR = 0.00027), Betaproteobacteria (4-22 %) (P = 0.13; FDR = 0.16) and Deltaproteobacteria (2-7 %) (P = 0.14; FDR = 0.17). Relative abundance of sequences representing members of class Actinobacteria (5-16 %) Acidobacteria subdivision Gp6 (1.8 – 7 %) and Acidobacteria subdivision Gp4 (1-3 %) was found to be significantly higher.

While at polluted sites the majority of the sequences were mapped for the members of class Bacteroidia (10-22%) (P = < 0.01; FDR = 0.00014), Anaerolineae (2-23%) (P = 0.041; FDR = 0.059), Clostridia (8-17%) (P = 0.00002; FDR = 0.00019), Synergistia (1-12%) (P = 0.00052; FDR = 0.001) Thermotogae (0.9- 15 %) (P = 0.0077; FDR = 0.014) and Flavobacteria (1.6 - 5%) (P = 0.5; FDR = 0.53). Classes affiliated with phylum Proteobacteria were annotated for Alphaproteobacteria (2.7-4.4 %) Betaproteobacteria (0.85-9.2 %), Gammaproteobacteria (6.5- 15 %) Epsilonproteobacteria (1-4.9 %) Deltaproteobacteria (9-14 %). The difference between relative abundance in the classes under Proteobacteria showed a reduction in the sequences annotated for Alpha, Beta and Gammaproteobacteria while a significant increase in the abundance was found to be in the members of class Epsilonproteobacteria and Deltaproteobacteria (Figure 2).

**Figure 2:**
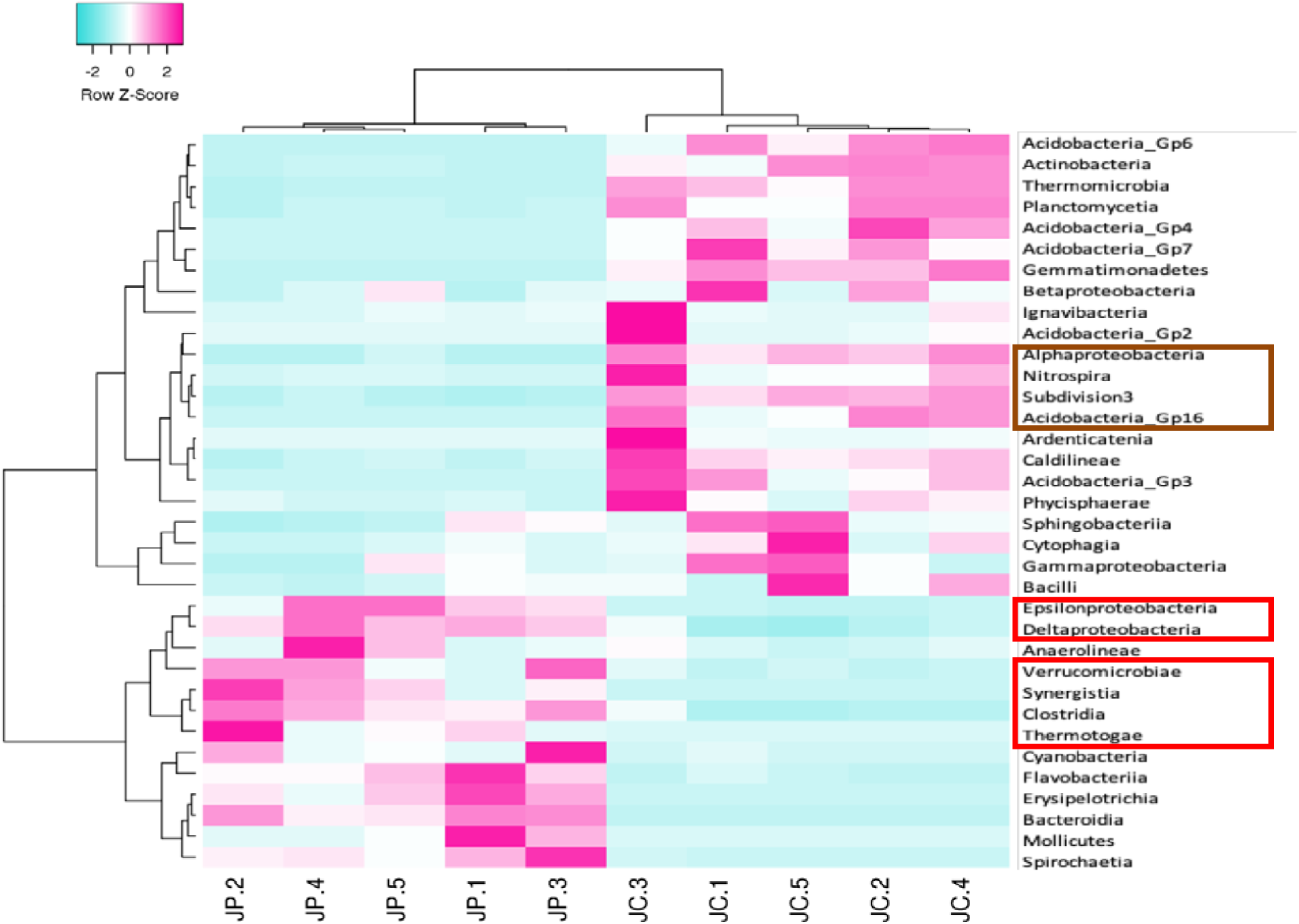
Heat map of differential abundance at Rank class (>1%) was mapped on RDP database. JP-1 to JP-5 represents polluted samples and JC-1 to JC-5 represents pristine/non-polluted samples. Z-score hierarchical clustering based on spearman rank correlation distance measurement method and complete linkage for linkage analysis. Each row represents one unique class representing more than 1 % abundance in each sample and each column represents one sample. The colour scale means the relative abundance, standard deviations from the mean, with blue for low abundance and pink for the high abundance values. Clustering was applied to both samples and class. Differential abundance at Rank class (>1%) was mapped on RDP database. JP-1 to JP-5 represents polluted soil sediments, JC-1 to JC-5 represents pristine/non-polluted soil samples.

The bacterial community structure of polluted and pristine sites of JIE was studied profoundly using amplicon sequence data at genera rank to understand the individual role and contribution of members in community assembly. Also this study offered a understanding the phenotypic and phylotypic shift in the community composition at polluted sites, as a result of amazing intelligence and interactions of microbial community at the perturbed site.

Strains of genera *Alcaligenes*, *Serratia* (only at JC-1 and JC-2), Gp6, *Gemmatimonas*, *Stenotrophomonas*, Gp4, Gp2, *Lacibacterium*, *Bellilnea*, *Pelagibus*, *Streptomyces*, *Bacillus*, *Thiohalophilus* and *Alcanivorax* were making around 29-48 % of the bacterial communities at pristine sites except at JC-3 where genera Gp3, Gp16 and *Longilinea* were also part of dominant group. These genera were drastically reduced at polluted sites making around 0.004-0.9 % of native community, except the genera *Alcaligenes* and *Serratia* were found to be present significantly higher at sites JP-4 and JP-5. This drastic reduction at abundance of these genera was found to be replaced by higher abundance of genera *Alkalitalea*, *Acholeplasma*, *Mariniphaga*, *Mesotoga*, *Lutaonella*, *Petrotoga*, *Desulfomicrobium*, *Aminivibrio*, *Macellibacteroides*.

*Sphaerochaeta, Levilinea, Smithella* and *Butyricoccus* making around 26-46 % of the total bacterial community at each site. These genera were found to be a part of rare microbiota at pristine sites contributing only 0.09-0.7 % in the community. About 60-76 % of the bacterial community of polluted sites was covered within top 20 bacterial genera, the relative abundance of many of these genera was slightly varied in among all 5 sites.

While at sites of pristine nature the distribution of top 20 genera was more subtle in numbers making around 45-69 % of the bacterial community, making the dominant community more widespread in diversity (Figure 3) The quest to find the core microbiome and the specific microbiome has led us to experience a unique pattern among all five sites of pristine and polluted environment which can be visualised in the Venn diagrams (Figure 4 and 5 respectively). On comparing the core microbiomes from both the environment significant differences were observed in the assembly of community players.

**Figure 3:**
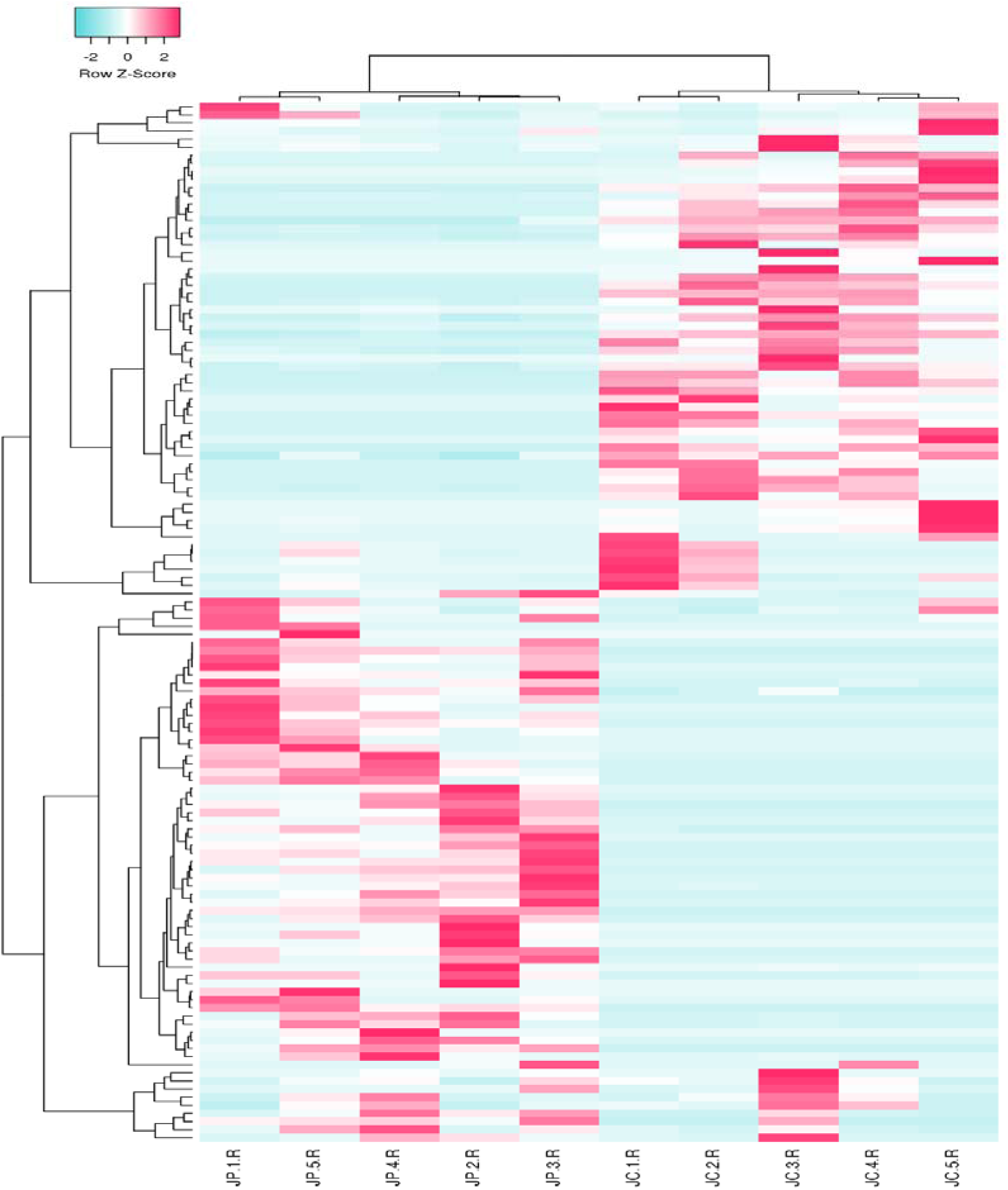
Heat map of differential abundance at Rank genera (>1%) was mapped on RDP database. JP-1 to JP-5 represents polluted samples and JC-1 to JC-5 represents control/pristine samples. Z-score hierarchical clustering based on spearman rank correlation distance measurement method and complete linkage for linkage analysis. Each row represents one genera unique genera from top 20 identified in each sample and each column represents one sample. The colour scale means the relative abundance standard deviations from the mean, with blue for low abundance and pink for the high abundance values. Clustering was applied to both samples and genera. JP- 1 to JP-5 represents polluted soil sediments, JC-1 to JC-5 represents pristine/non-polluted soil samples.

**Figure 4:**
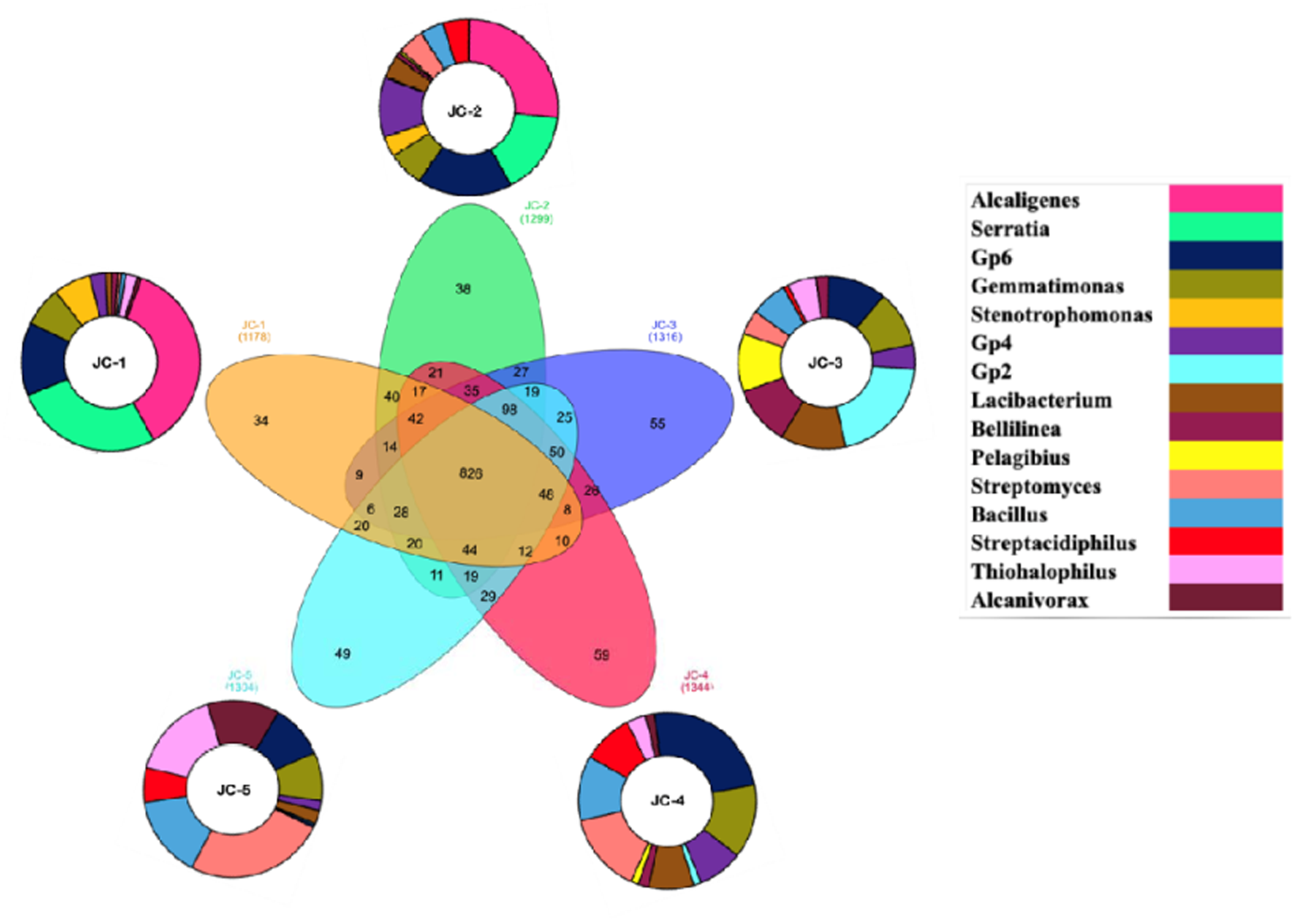
Integrated diagram at rank genera showing the number of core and unique genera annotated from each of the pristine sites of JIE in the Venn diagram at centre while the differential abundance of top 5 genera annotated from each pristine/non-polluted site is shown in surrounding doughnut charts. For each unique genera a unique colour was assigned. The size of each section represents the percentage abundance of a single genera in dominant community structure.

**Figure 5:**
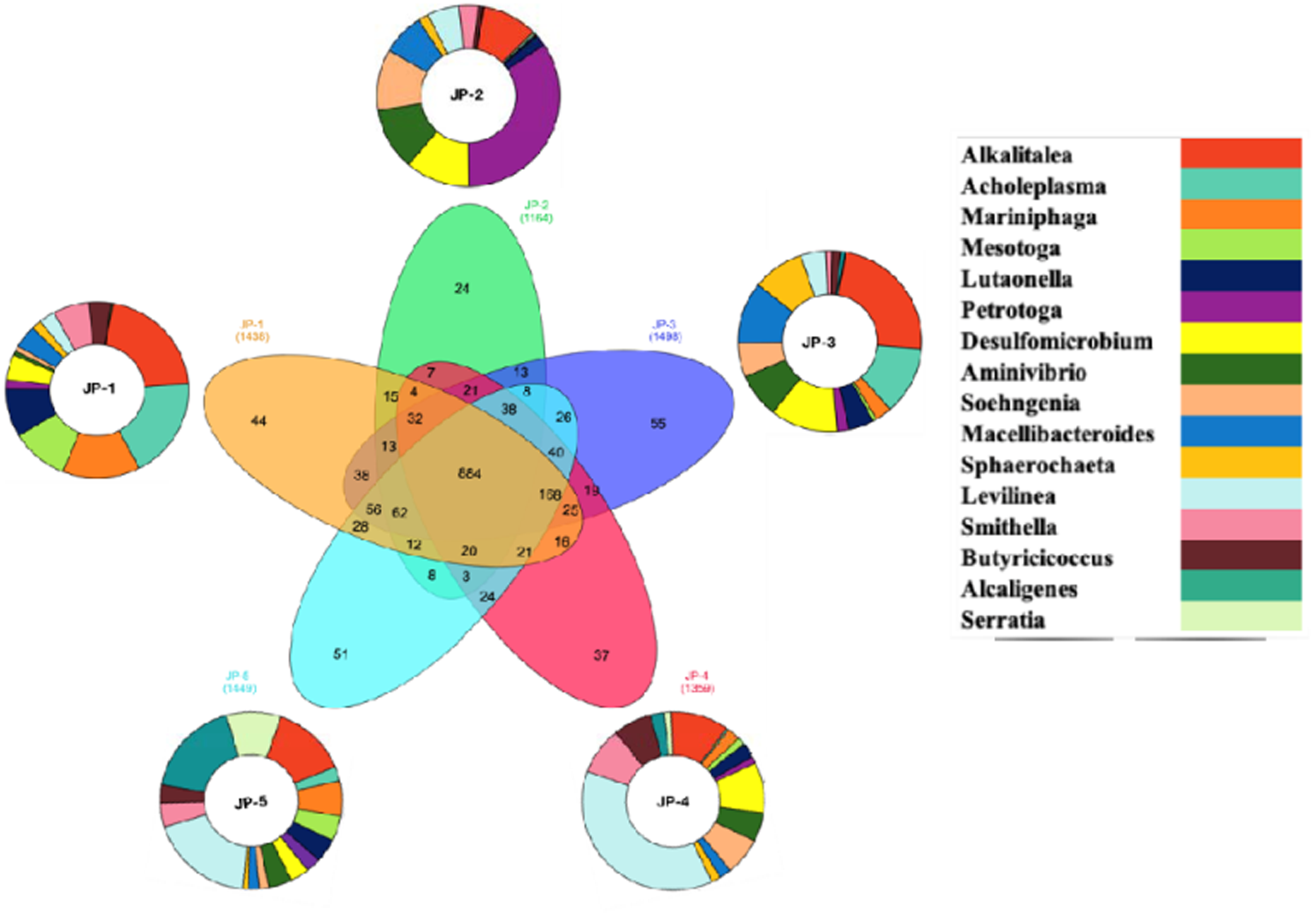
Integrated diagram at rank genera showing the number of core and unique genera annotated from each of the polluted sites of JIE in the Venn diagram at centre while the differential abundance of top 5 genera annotated from each polluted site is shown in surrounding doughnut charts. For each unique genera a unique colour was assigned. The size of each section represents the percentage abundance of a single genera in dominant community structure.

#### 3.2.3 Archaeal Community Structure of Jetpur Soil

The analysis of 16S rRNA gene indicated that both domains Bacteria and Archaea were detected in all samples. Although most archaeal phyla were present at low abundance (less than 1% of the total microbial abundance), the dominant archaeal phylum was Euryarchaeota at all 10 sites. The archaeal community was found to be more diverse at pristine sites when compared to that of polluted sites when mapped at rank Phylum. A significant reduction of the members of phylum Pacearchaeota, Aenigmarchaeota, Parvarchaeota, Thaumarchaeota (ammonia oxidizers) was seen at all five polluted sites (JP-1 to JP-5) which were showing abundance in range of 5 to 36 % at pristine sites (JC-1 to JC-5). While members of Euryarchaeota alone were dominating at polluted sites by ranging from about 78 to 99% in abundance (Figure 6).

**Figure 6:**
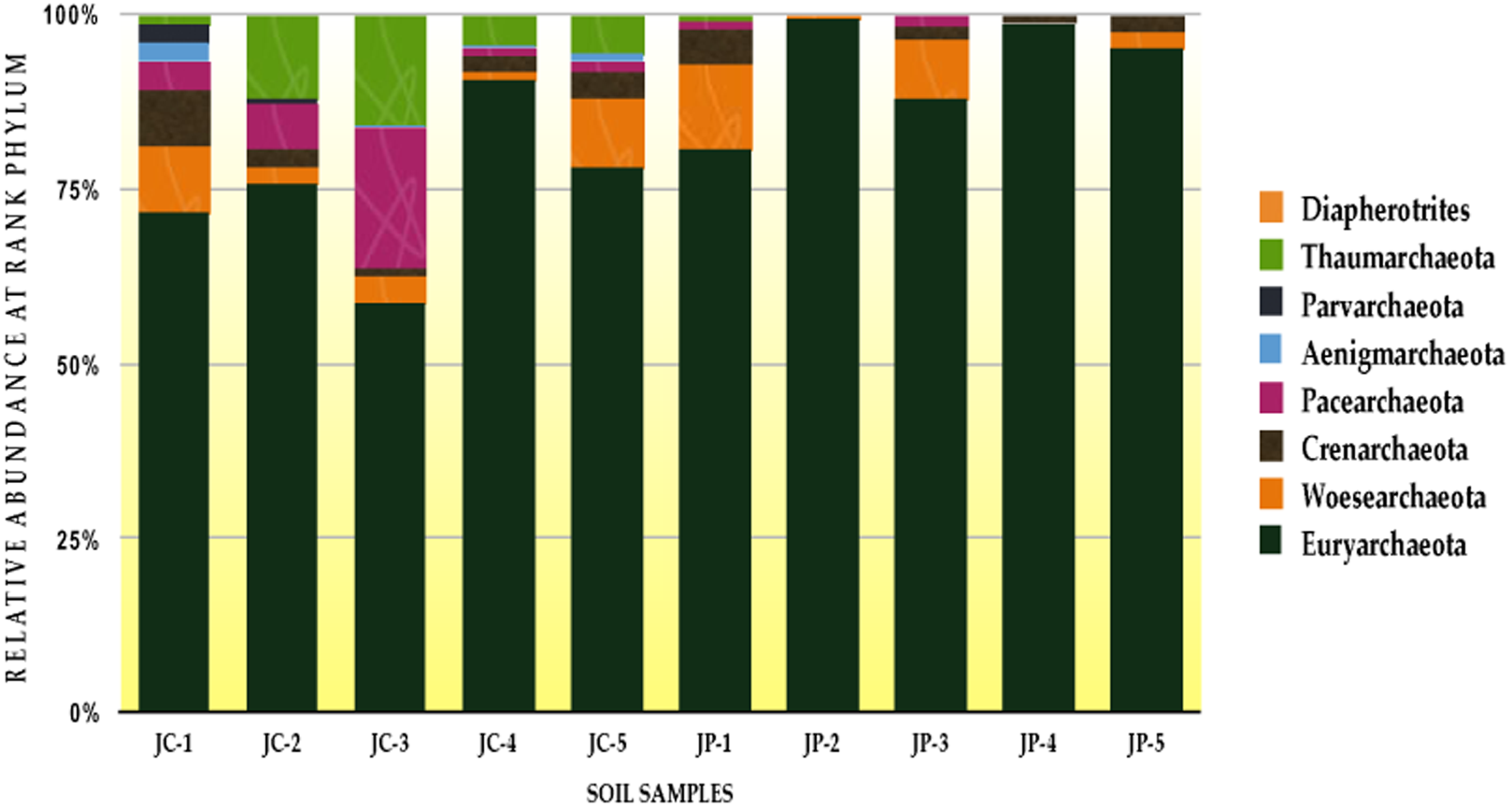
Differential abundance of Archaeal community depicted at rank phylum for 10 sites of JIE (mapped on RDP database). JP-1 to JP-5 represents polluted soil samples, JC-1 to JC-5 represents pristine/non-polluted samples.

At genera level a complete shift was observed between pristine and polluted microbial community. The genera belonging to class *Methanomicrobia* (53-95 %) were making the most of the archaeal community at polluted sites (JP-1 to JP-5) including *Methanosaeta* (11-69 %) *Methanocalculus* (3.5– 52.5 %), *Methanospirillum* (2.8 – 25 %), *Methanocorpusculum* (0.4– 24.7 %) *Methanogenium* (0-2.6 %) *Methanoplanus* (0 -3.8 %). Genera *Halomarina* (0- 8.9%) and *Thermosphaera* (0-3.3 %) respectively from class Halobacteria (0-9 %) and Thermoprotei (0-3 %) were also found to be differentially abundant in polluted sites. While pristine sites (JC-1 to JP-5) were found to be dominated by members of an entirely different class (Thermoplasmata) of phylum Euryarchaeota. *Methanomasiliiicoccus* was dominating the pristine sites with relative abundance ranging from 56 to 87 % (Figure 7).

**Figure 7:**
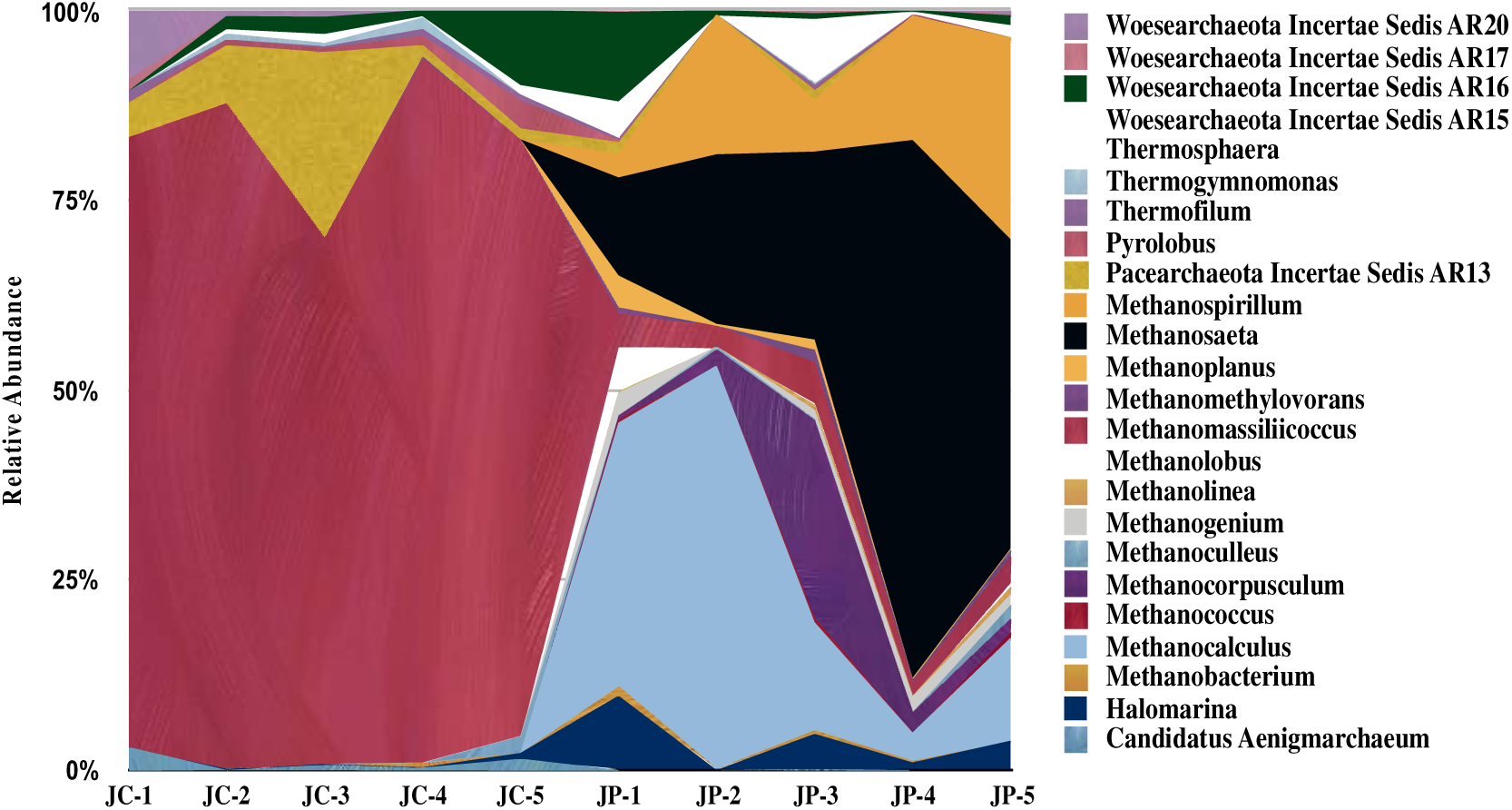
Representation of differences between relative abundance of archaeal genera from polluted and pristine site amplicon sequences annotated on RDP database showing genera showing a minimum of 1 percentage abundance on each of the 10 sites. JP-1 to JP-5 represents polluted soil sediments, JC-1 to JC-5 represents pristine/non-polluted soil samples.

### 3.3 Statistical analysis of diversity among samples

The community changes between polluted and pristine habitats were analyzed using different diversity indices. Diversity is the measure of both taxon richness and evenness. Alpha diversity estimation indicated a little diversity variation within the samples, but the bacterial community was formed by great bacterial diversity at polluted sites (Table 3). Through Beta-diversity estimation (Figure 8) the induced shift in bacterial community (both taxonomically and phylogenetically) at polluted ecosystem of JIE was clearly observed. The variations among the ten bacterial communities (five each from polluted and pristine) were grouped with a total variance of 94 %, according to the effect of industrial activities.

**Figure 8:**
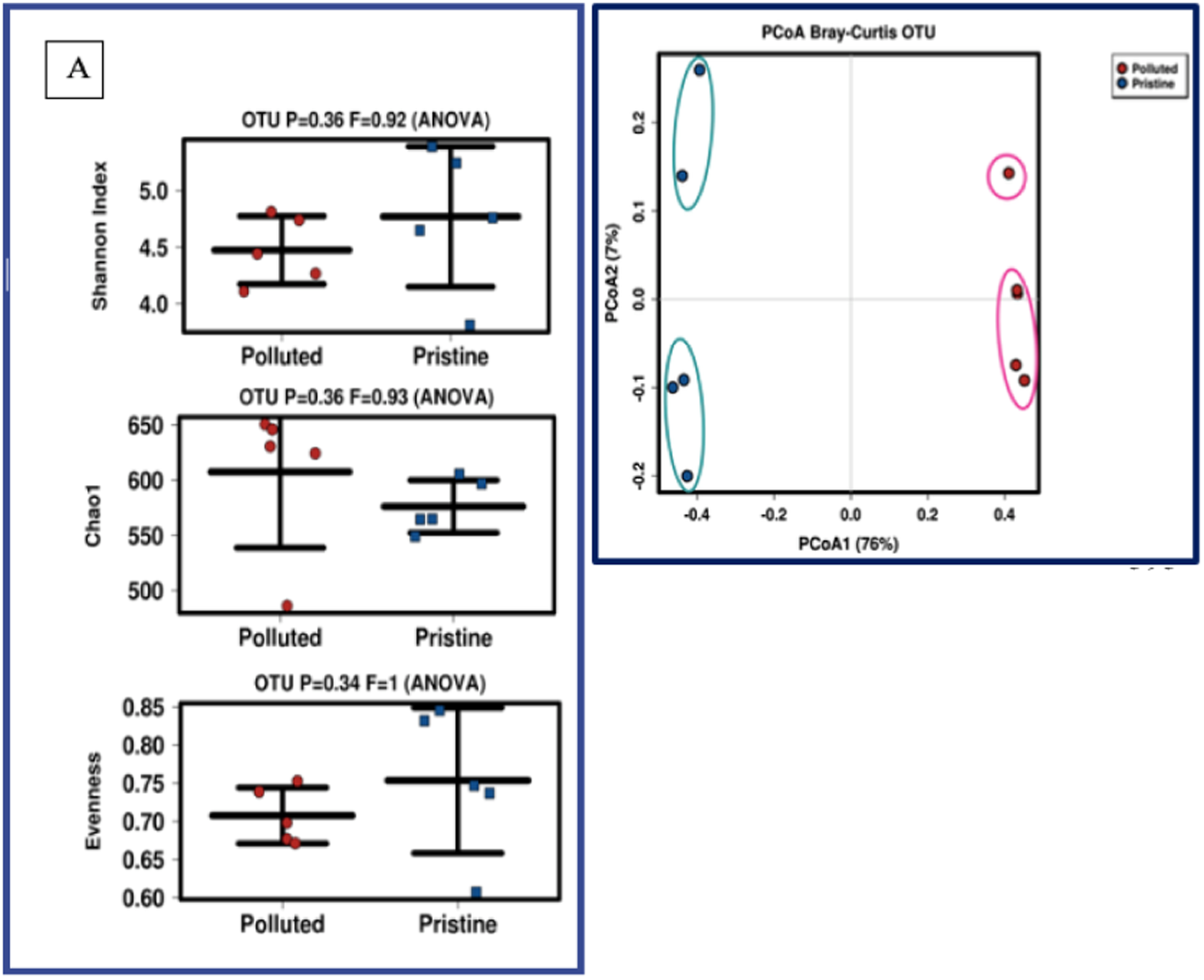
Representation of differences between relative abundance of archaeal genera from polluted and pristine/non-polluted site amplicon sequences annotated on RDP database showing genera showing a minimum of 1 percentage abundance on each of the 10 sites. JP-1 to JP-5 represents polluted sediments, JC-1 to JC-5 represents polluted and non-polluted samples.

**Table 3:**
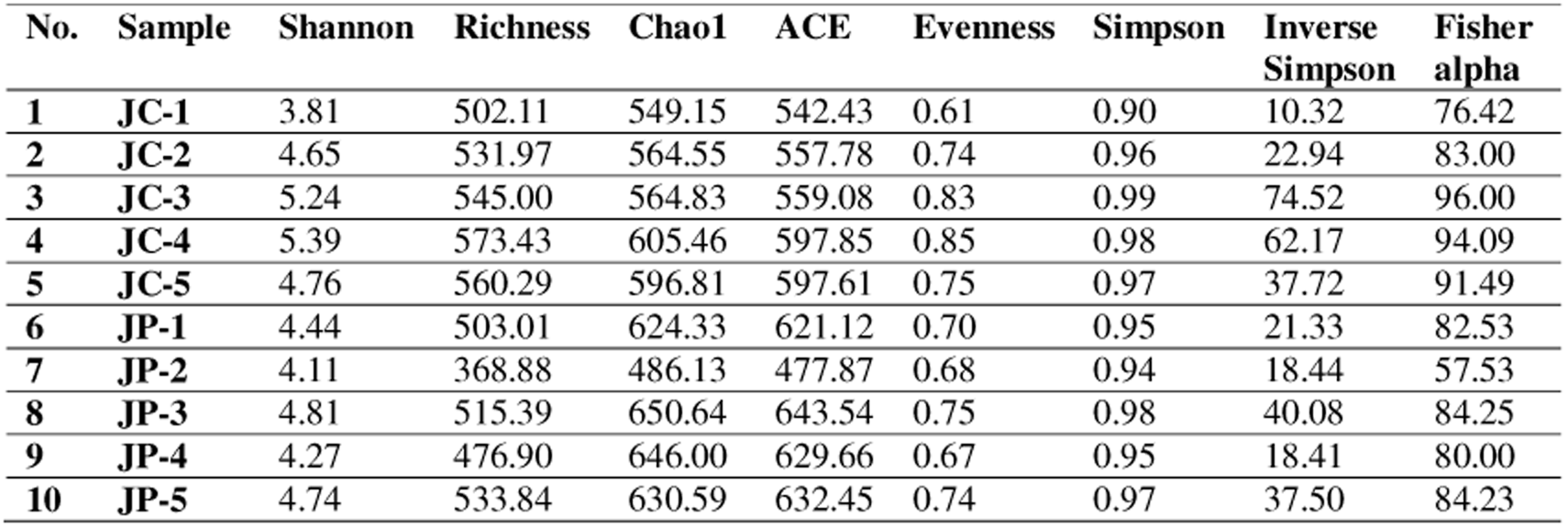
Alpha diversity measures estimated for polluted (JP-1 to JP-5) and pristine/non-polluted (JC-1 to JC-5) sites of Jetpur Industrial Estate, Jetpur.

### 3.4 Predictive (microbial) metaphenomics: In-silico approach for system microbiology

Soil microbiomes usually operate in dynamic social networks where populations and consortia are shaped by continuous cooperative and competitive interactions to maintain their ecological niches. To understand the microbial (here bacterial and archaeal) response as a system adapting to anthropogenic perturbations, assessing the phenotypic (i.e. a metaphenome) traits of autochthonous communities in polluted environments is essential.

The results from amputed metegenomic based on 16S rRNA gene sequencing and using trait-coverage estimation, the organismsm-level and/or system-level microbiome phenotype were predicted. Functional trains were predicted using PiCRUST, where 303 distinct metabolic pathways from polluted samples were annotated from KEGG/KO ontology (a three tier classification system) defined by the BRITE hierarchy. Similarly for non-polluted habitats, functions related to 306 pathways were mapped. These pathways are coupled with both eukaryotic and prokaryotic systems. The non-polluted ecosystem was shown to have more genes associated with eukaryotes. These pathways were then grouped into secondary and tertiary organisational tier to further define the subtle differences in the functionality between the both microbial communities. Apart from this organism level and system level interactive prediction based analysis, the phenomic features of the communities were studied to reconstruct a “Metaphenome” analysis from both communities.

The predictive features of polluted (as well as non-polluted microbiome) indicated the high abundance of genes and proteins associated with metabolism of xenobiotic compounds, besides the functions for nitrogen, carbon, lipids, energy, amino acids metabolism, membrane transport, etc. were also observed. A significant increase was observed in the community of polluted ecosystem in metabolic capacities for acetoclastic methanogenesis, aerobic anoxygenic phototrophy, reductive acetogenesis, respiration of sulfur compounds, dark sulphite oxidation and various fermentative processes. Several catabolic pathways for degradation of complex aromatic compounds were annotated from polluted ecosystem. An increase was observed for aromatic and oil biodegradation capabilities (Figure 9).

**Figure 9:**
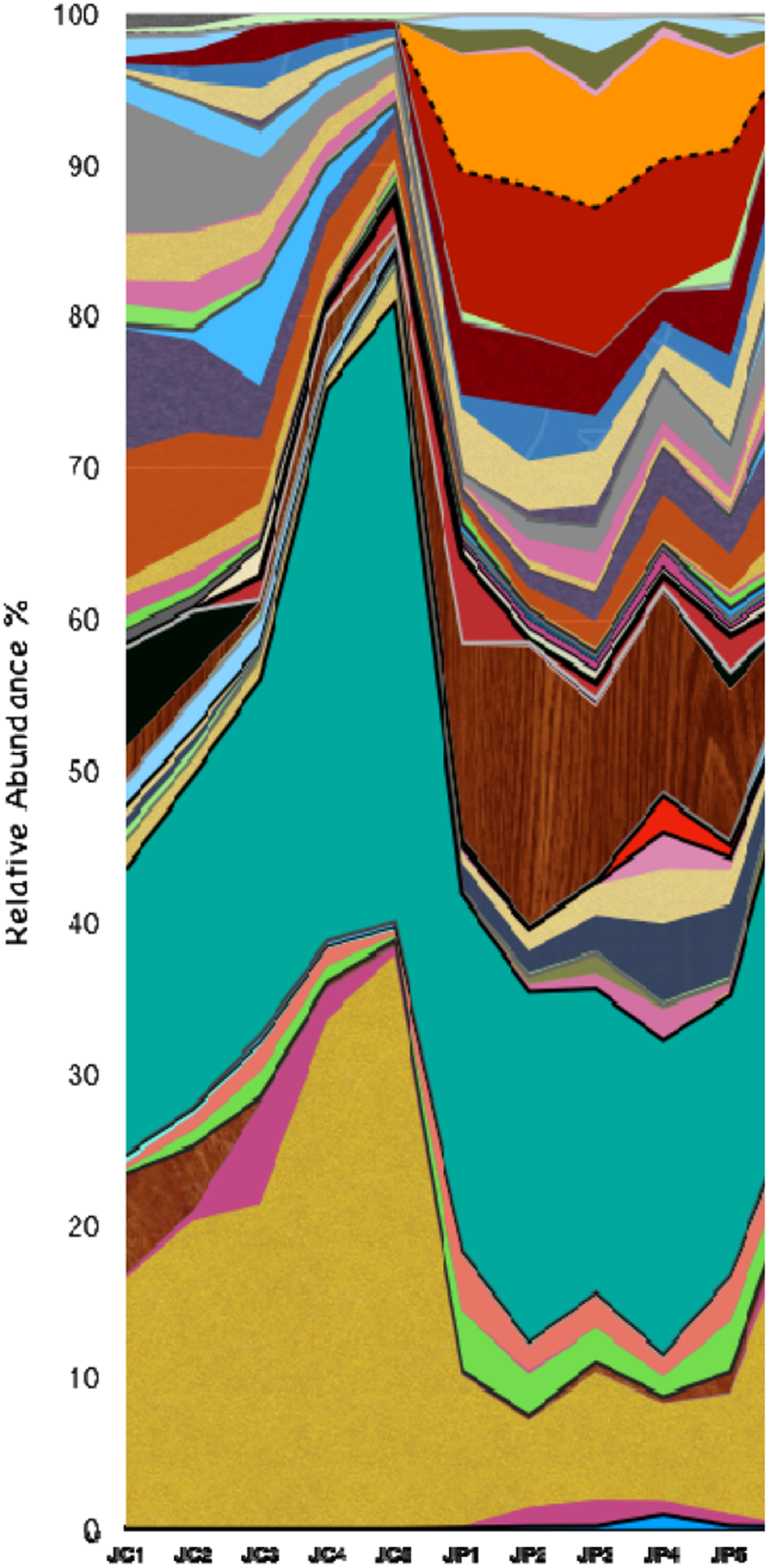
Metaphenomic characteristics of polluted and pristine communities mapped for the sites of Jetpur Industrial estate, Jetpur, Gujarat. The predicted metaphenome for all 10 sites of JIE can be read from downside upward direction from acetoclastic, methanogenesis to xylanolysis.

It is worthy to mention that the NSTI: Nearest Sequenced Taxon Index, scores for predicted metagenomes ranges from 0.12 to 0.14 for polluted metagenome, with mean of 0.19 and that for non-polluted it ranges from 0.14 to 0.21 and a mean of 0.17. While the NSTI score for overall prediction were between 0.14 and 0.22 with overall mean of 0.18, found to have fairly accuracy for soil samples (Robinson et al., 2016, Langille et al., 2013). Based on these findings a diagrammatic representation was created representing top 5 genera from each of the polluted and non-polluted sites in which each phenotype was represented as a unique coloured square box and various phenotype for a single genera were added together to claim their overall characteristics in a comparative manner (Figure 10).

**Figure 10:**
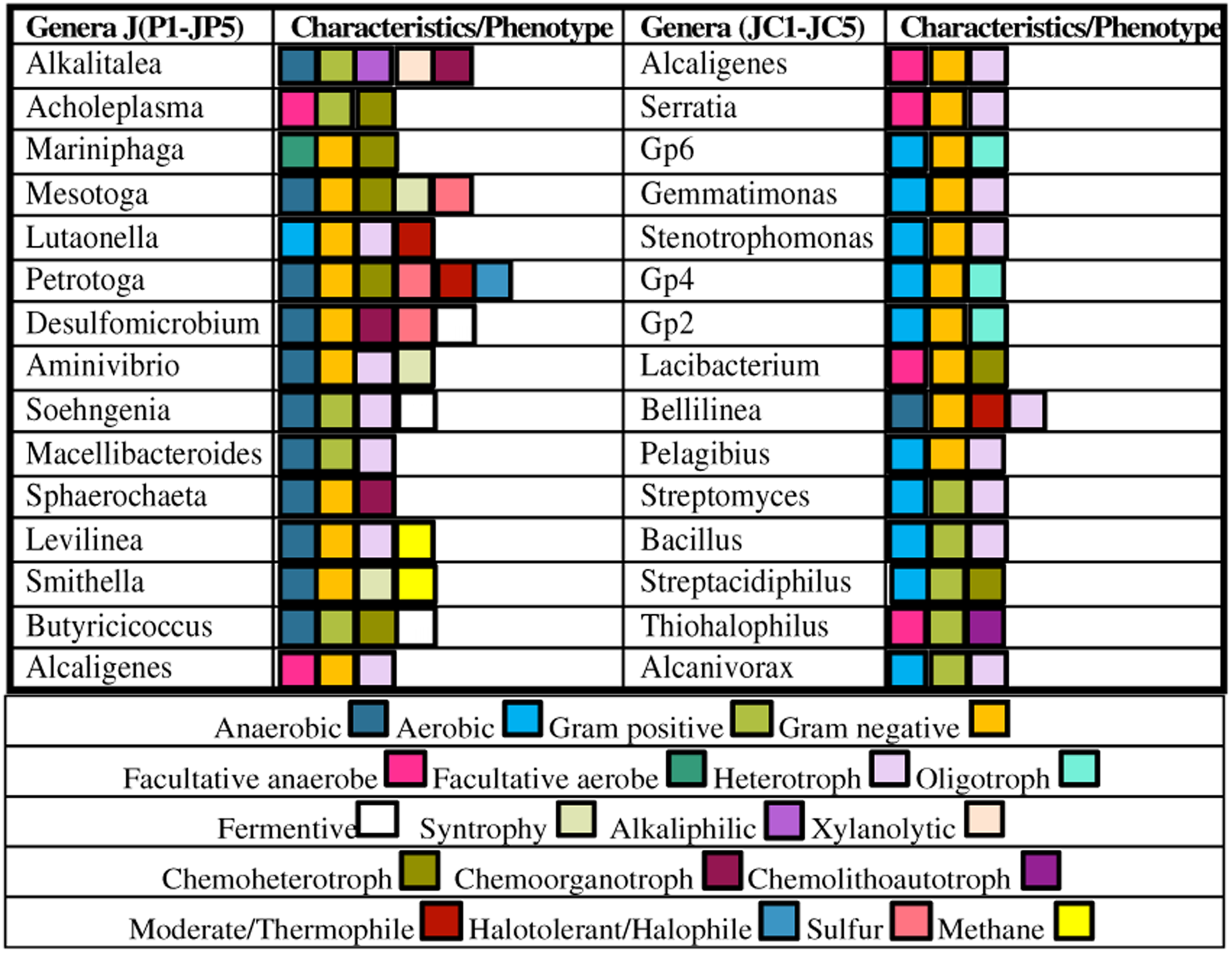
Diagrammatic representation for various phenotypic characteristic for 5 most dominant genera from polluted and pristine ecosystem of JIE. Each coloured block represents a unique phenotype. The more the number of blocks the more characters are affiliated to a particular genera.

The community interactions and co-occurrence patterns for maintaining the microbial ecology of both polluted and non-polluted sites were identified using network analysis. The predictive analysis revealed the co-occurrence and strong interactions among members of phylum Proteobacteria, Actinobacteria, Firmicutes, Acidobacteria, Planctomycetes, Verrucomicrobia, Nitrospirae and Cyanobacteria at non-polluted pristine sites of JIE depicting the normal soil microbiome. While at polluted site very strong interactions and co-occurrence patterns were observed for species of phylum Thermotogae, Synergistetes, Euryarchaeota, Bacteroidetes and Fusobacteria indicating a selective ecological adaptation (microbial intelligent) of the previously undisturbed community, shifting towards diverse fermentative, anaerobic, and sulfur respiratory pathways (Figure 11 and Figure 12).

**Figure 11:**
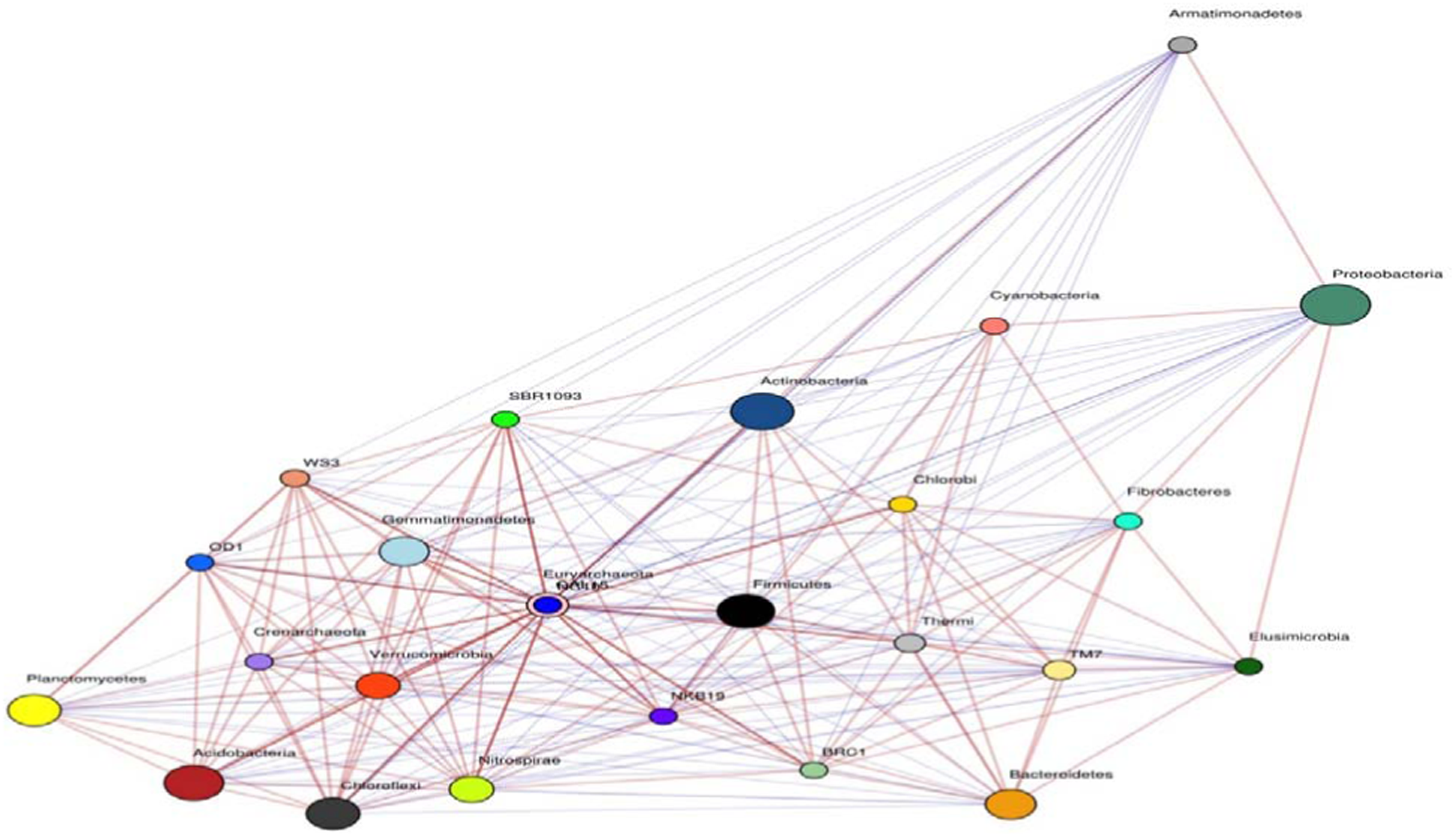
The community interactions and co-occurrence patterns of pristine/non-polluted microbial ecosystem at JIE. The node size is representative for the abundance of that particular phylum, while the blue edges represents negative co-occurrence and interactions while the yellow represents positive correlation between the nodes.

**Figure 12:**
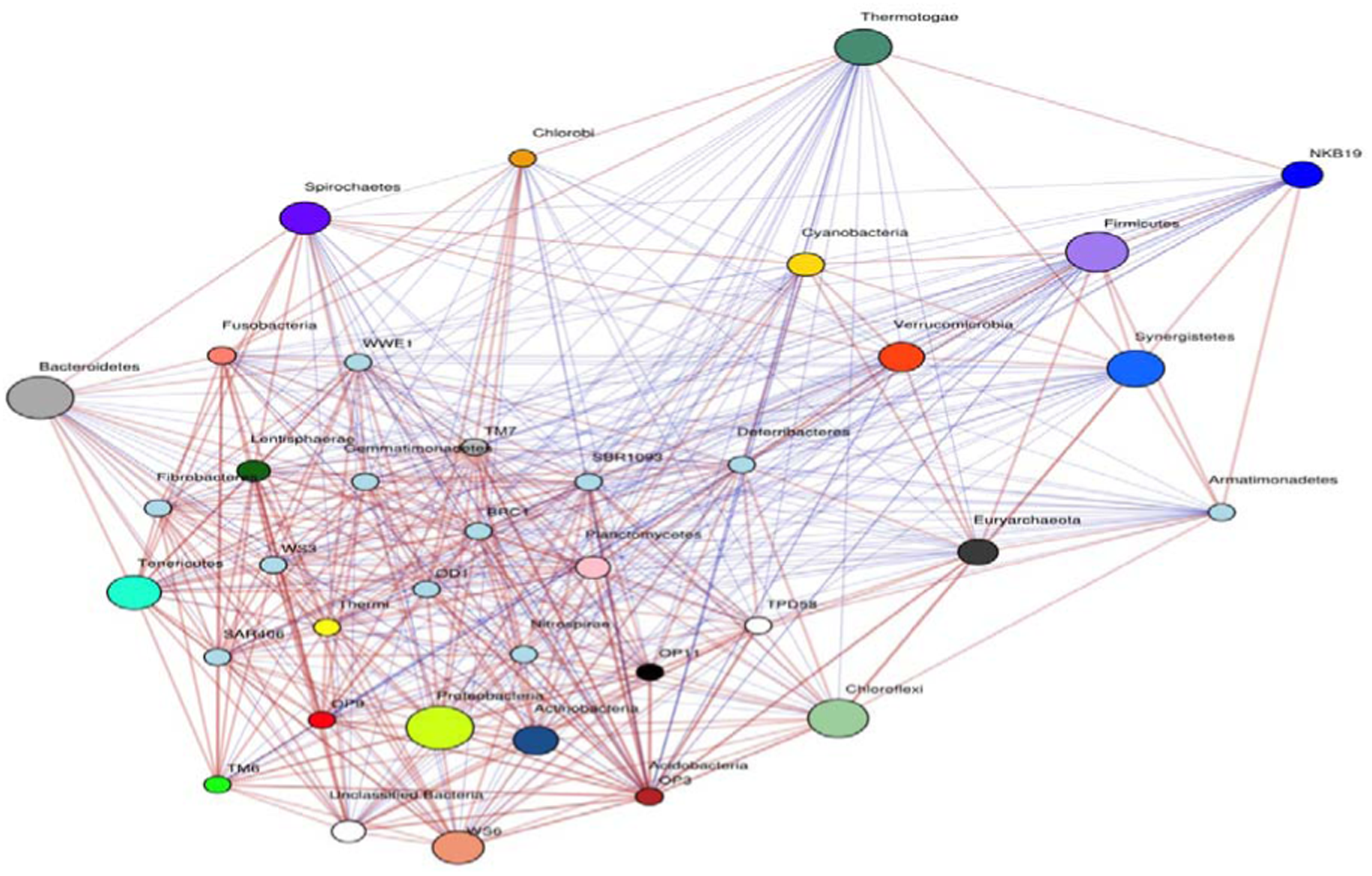
The community interactions and co-occurrence patterns of polluted microbial ecosystem at JIE. The node size is representative for the abundance of that particular phylum, while the blue edges represents negative co-occurrence and interactions while the yellow represents positive correlation between the nodes.

### 3.5 Functional Potential of Polluted Microbiome of JIE using shot-gun metagenomics

#### 3.5.1 Sulphur Metabolism

Like carbon and nitrogen, sulphur is also an important element for life. Besides its structural relevance in protein biology, it has a significant role in central biochemistry (in all forms of life) as a carbon carrier and stable redox centre (Klotz et al., 2011). Sulphur is the sixth most abundant element in the microbial biomass; which has a profound effect on biosphere as well as in geochemistry/bio-geochemistry (Klotz et al., 2011; Plugge et al., 2011). A complex sulphur cycle exists in the biosphere depending upon the inter-changing valence of sulphur atom (–2 (sulphide) to +6 (sulphate)). Since many of the sulphur reactions are non-spontaneous (under ambient conditions), prokaryotic metabolism is imperative for recycling of elemental sulphur in the biosphere (Klotz et al., 2011). Moreover, in the absence of molecular oxygen and nitrate, microorganisms prefer sulphur compounds either as electron donors or electron acceptors for energy transformation/conserving metabolism (Purcell et al., 2014; Plugge et al., 2011; Yamamoto et al., 2011; Klotz et al., 2011).

With the above facts, a similar bi-functional (oxidation-reduction reactions) properties of autochthonous microorganisms of JIE were observed for their sulfur metabolic capabilities. The functional capabilities for sulphur metabolism of autochthonous microorganisms of JIE were characterized into ten distinct pathways and can be broadly clustered as observed in Figure 13. The autochthonous microbial community of polluted habitat of JIE has a significant inherent potential for sulphur metabolism. The phenotype may be expressed for alternate mode of respiration using sulphate, sulfite, thiosulfate and sulphur compounds. Since the polluted habitat of JIE had characteristic anaerobic and anoxic micro-niches, this energy conserving metabolism was catalysed though dissimilatory sulfur reduction pathways.

**Figure 13:**
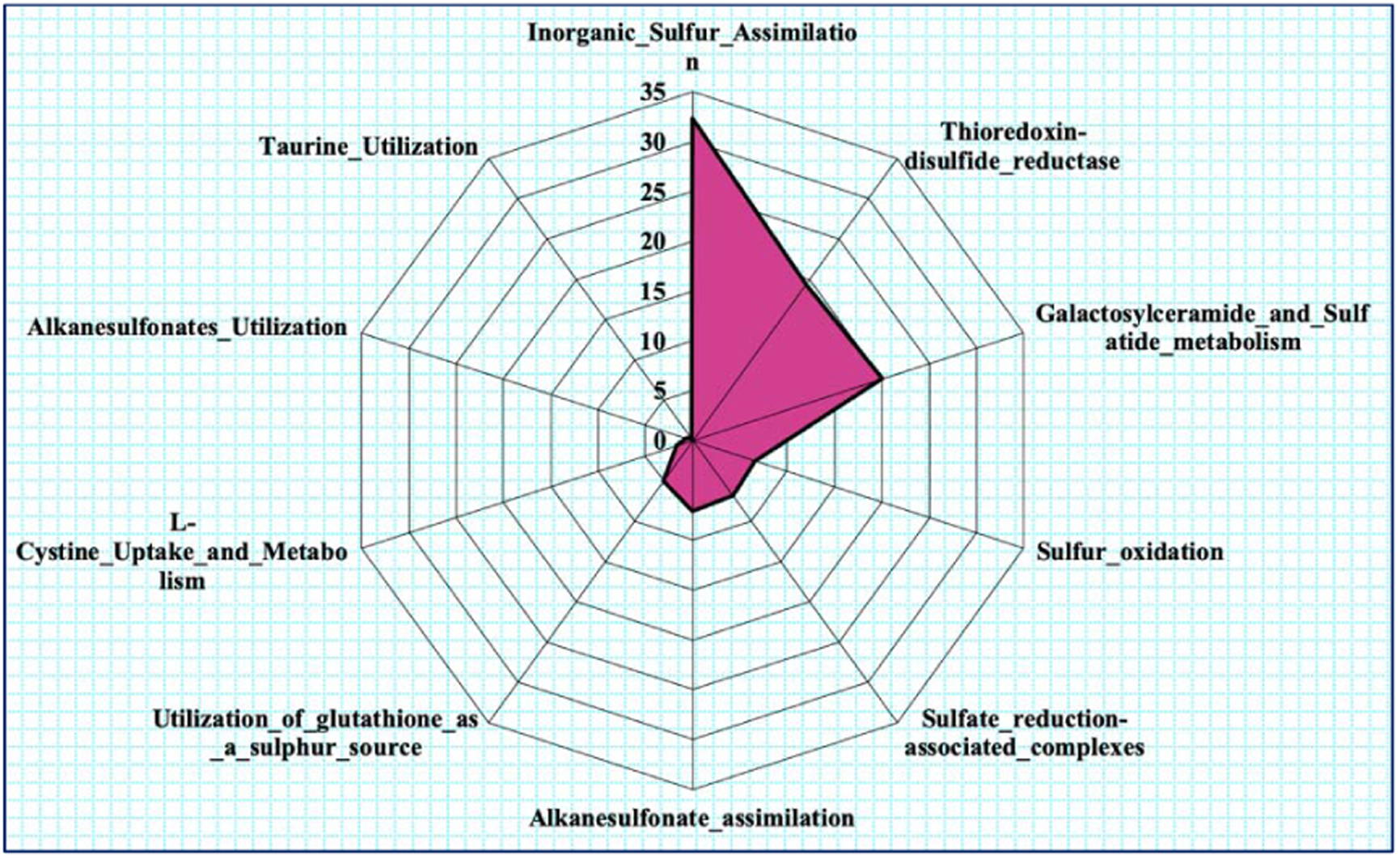
Relative differential abundance of functional features for various pathways for metabolism of Sulfur (Level-2), annotated from metagenome of polluted soils of Jetpur Industrial Estate, Jetpur. The functions were mapped on Subsystem database on MG-RAST server with default parameters.

For both, reduction and oxidation in S-loop (sulphur cycling), 11 gene families are involved and atleast representative genes and proteins from each families were annotated. During dissimilatory sulphate reduction (anaerobic/anoxic sulphur respiration), where sulphate is reduced into hydrogen sulphide (H_2_S), genes encoding for reversible adenylylsulfate reductase AB (*aprAB*) [EC: 1.8.99.2], reversible sulphate adenylyltransferase (*sat*) [EC: 2.7.7.4] and dissimilatory (bi)sulfite reductase (*dsrAB*), [EC: 1.8.99.5] were annotated from polluted habitat of JIE. For assimilatory sulphate reduction, besides *sat*, genes encoding for sulfite reductase (*cysJI*), bifunctional enzyme cysN / cys C (*cysNC*) (sulphate adenylyltransferase and adenylylsulphate kinase) and phosphoadenosine phosphosulfate reductase (*cysH*) [EC: 1.8.4.8; 1.8.4.10] were annotated. For sulphur and thiosulphate oxidation (thiosulphate to sulphate), sulphite dehydrogenase (quinone) (*soeABC*) [EC: 1.8.5.6] and L-cysteine S-thiosulfotransferase (*soxA*) [EC: 2.8.5.2] were annotated. For inorganic sulphur transformations, thiosulphate reductase / polysulfide reductase (*phsABC*) [EC: 1.8.5.5], sulphydrogenase / dehydrogenase I (*hydA*) [EC: 1.12.1.3; 1.12.1.5], *hydGB* [EC:1.12.98.4] were annotated.

In sulphate-sulphur, genes for assimilation of sulphate/thiosulphate transport system ATPbinding protein (*cysA*) [EC: 7.3.2.3], sulphate/thiosulphate transport system permease protein (*cysUW*) were annotated. In organic sulphur transformations, alkanesulphonate monooxygenase (*ssuD*) [EC: 1.14.14.5], taurine transport system substrate-binding protein (*tauAC*), taurine dioxygenase (*tauD*) [EC: 1.14.11.17] amongst the other genes encoding the enzymes for sulphur metabolism were annotated.

Genes encoding the enzymes and proteins for various sulphur metabolisms were phylogenetically assigned to members of order Desulfovibrionales, Chromatiales, Pseudomonadales, Rhodobacterales, Rhodocyclales, Eurotiales, Thermotogales, Clostridiales.

The genes were taxonomically assigned to *Desulfomicrobium* spp., *Allochromatium* spp., *Desulfovibrio* spp., *Pseudomonas* spp., *Rhodobacter* spp., *Thauera* spp., *Thioflavicoccus* spp., *Desulfohalobium* spp., *Azotobacter* spp., *Clostrridium* spp., *Petrotoga* spp., *Sulfuricella* spp. and many other significantly less contributing species.

#### 3.5.2 Nitrogen Metabolism

Nitrogen is an essential element for the all forms of life. The nitrogen cycling in the polluted environment is one key function, because at perturbed ecosystem nutrients are often limited due to an unbalanced C:N ratio (Yergeau et al., 2014). Moreover, nitrogen metabolism is also important in the habitats/ecosystem with limited available oxygen concentration, where microorganisms prefer nitrate/nitrogen compounds either as electron donors or electron acceptors for energy transformation and conserving metabolism (Erisman et al., 2007).

Most part of the nitrogen cycle (N-loop) in the environment is governed by the microorganisms (Ward and Jensen, 2014; Meyer, 1994). With the current state of our understanding, microbial transformations of nitrogen consist of six (dissimilatory nitrogen reduction to ammonia, assimilatory nitrogen reduction, N_2_ fixation, denitrification, ammonification and anaerobic ammonia oxidation) interrelated but distinct processes (cycle) proceed in a defined manner, where nitrogen in organic or inorganic compounds changes its redox states between –3 and +5 (Thomson et al., 2012; Scott et al., 2014; Nelson et al., 2016).

For N-loop, 18 gene families are involved (Zumft et al., 1997; Meyer, 1994; Thomson et al., 2012) and at lease representative genes and proteins from each family were annotated from JIE. In dissimilatory nitrate reduction, where nitrate is concerted into ammonia, genes encoding for nitrite reductase (*nirBD*) [EC: 1.7.1.15], nitrate reductase/nitrite oxidoreductase (*narGH*) [EC: 1.7.5.1; 1.7.99.-] were annotated. During denitrification, where nitrate is converted into nitrogen, nitric oxide reductase (*norBC*) [EC: 1.7.2.5], nitrite reductase (NO forming)/hydroxylamine reductase (*nirS*) [EC: 1.7.2.1; 1.7.99.1], periplamic nitrate reductase (*napA*) [EC: 1.7.99.-] were annotated. For nitrate assimilation genes encoding the nitrate/nitrite transport system substrate-binding protein/permease protein/ATP-binding protein (*nrtABC*) [EC: 3.6.3.-] were annotated. The genes categorized as subsystem level 2 for nitrogen metabolism are represented in Figure 14.

**Figure 14:**
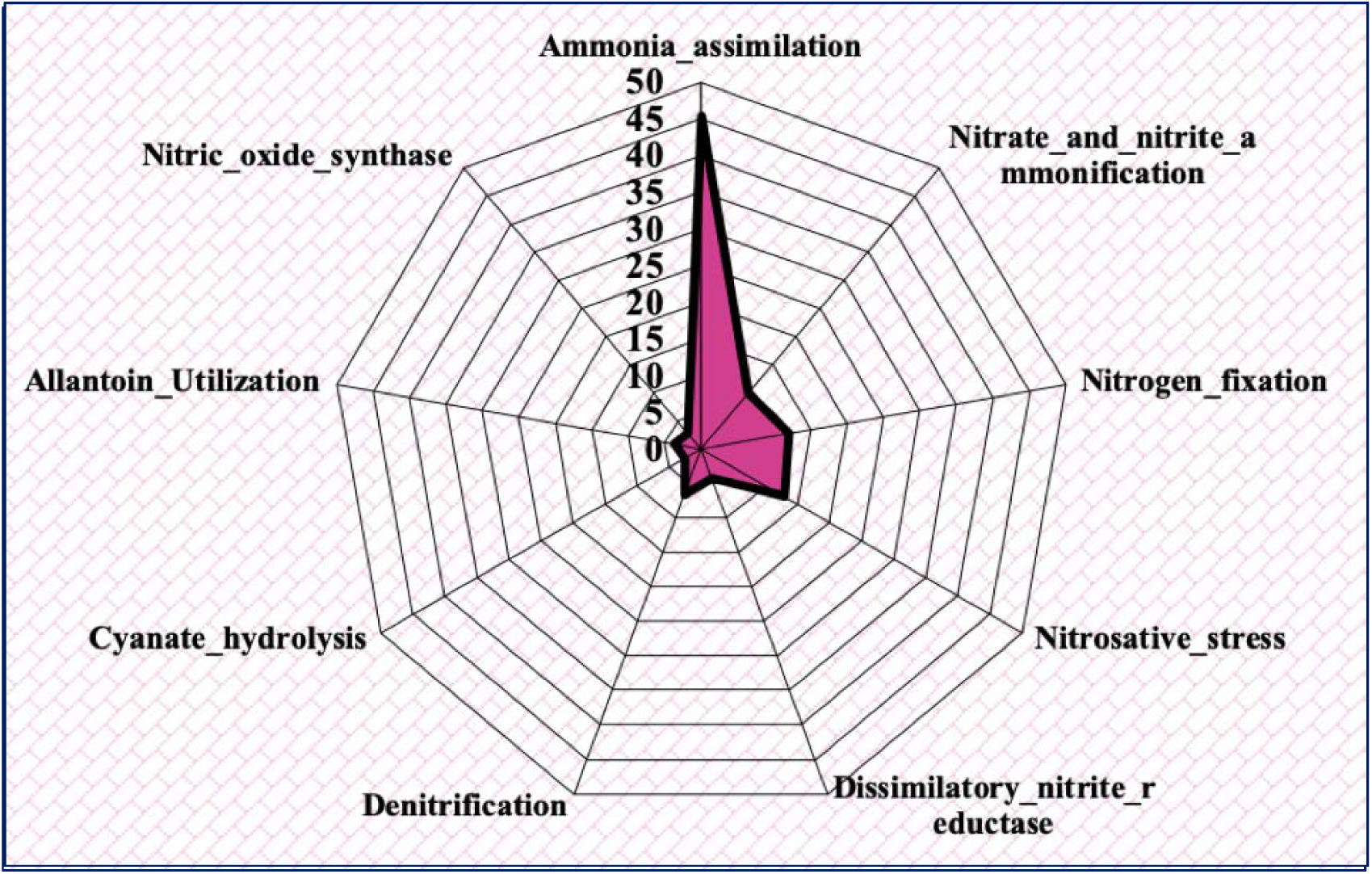
Relative differential abundance of functional features for various pathways for metabolism of Nitrogen (Level-2), annotated from metagenome of polluted soils of JIE. The functions were mapped on Subsystem database on MG-RAST server with default parameters.

Genes encoding the enzymes and proteins for various nitrogen metabolisms were phylogenetically assigned to Thermotogales, Sphingobacteriales, Methanosarcinales, Rhodocyclales, Spirochaetales, Burkholderiales, Chromatiales, Desulfovibrionales, Synergistales, Rhodobacterales, Desulfobacterales, Oscillatoriales, Rhodospirillales, Campylobacterales, Pseudomonadales, Clostridiales, Bacteroidales, Desulfuromonadales, Xanthomonadales and many more. They are taxonomically identified from species of genera *Petrotoga, Chitinophaga, Methanosaeta, Thauera, Sphaerochaeta, Mesotoga, Desulfomicrobium, Sorangiineae, Rhodobacter, Methanospirillum, Allochromatium, Arthrospira, Ralstonia, Desulfobulbus, Achromobacter, Azospirillum, Anaerobaculum, Arcobacter, Aminobacterium* and *Desulfovibrio* mainly.

#### 3.5.3 Methane Metabolism (Methanogenesis)

Although methanogenesis was thought be a niche specific process of (obligatory) anaerobic ecosystem, the polluted metagenome of anoxic ecosystem of JIE provided strong evidence of methanogenic potential of autochthonous microbial community. A *mcrA* gene encoding methyl-coenzyme M reductase [EC 2.8.4.1], an important gene catalyzes the final reaction in biogenic methanogenesis was annotated. Along with *mcrA*, *mcrBG* as well as gene encoding the enzyme tetrahydromethanopterin S-methyltransferase [EC:2.1.1.86] (*mtrBCDEH*) were annotated. The annotation of hydrogenase (*viz.* F420-non-reducing hydrogenase), and genes involved in pathways for acetyl-CoA and methylamine metabolism, evidently indicates the co-existence of hydrogenotrophic, acetoclastic and methylotrophic methanogenesis at polluted ecosystem of JIE.

Further, gene encoding heterodisulfide reductase (hdr) [EC:1.8.7.3; 1.8.98.4; 1.8.98.5; 1.8.98.6] was also annotated, suggesting the intracellular transport associated with methane metabolism (Reid et al., 2018). It also utilizes F420-non-reducing hydrogenase (*mvhADG*) to complete H_2_-dependent reduction of heterodisulfide to ferredoxin and was annotate from polluted ecosystem (Reid et al., 2018). Gene encoding the enzyme F420-dependent N5, N10-methylenetetrahydromethanopterin reductase [EC 1.5.99.11] was also annotated. At polluted site of JIE majority of the players performing the functions for methane metabolism were annotated from archaeal phylum Euryarchaeota along with from members of phylum Thermotogae.

#### 3.5.4 Stress Response

The annotation for stress response mechanism revealed that the genes for oxidative stress response was the most abundant one among other subsystem level 2 categories. Along with this genes for osmotic stress, detoxification, heat shock and cold shock responses were also annotated. A detailed description of functions under this category can be understood in Figure 15.

**Figure 15:**
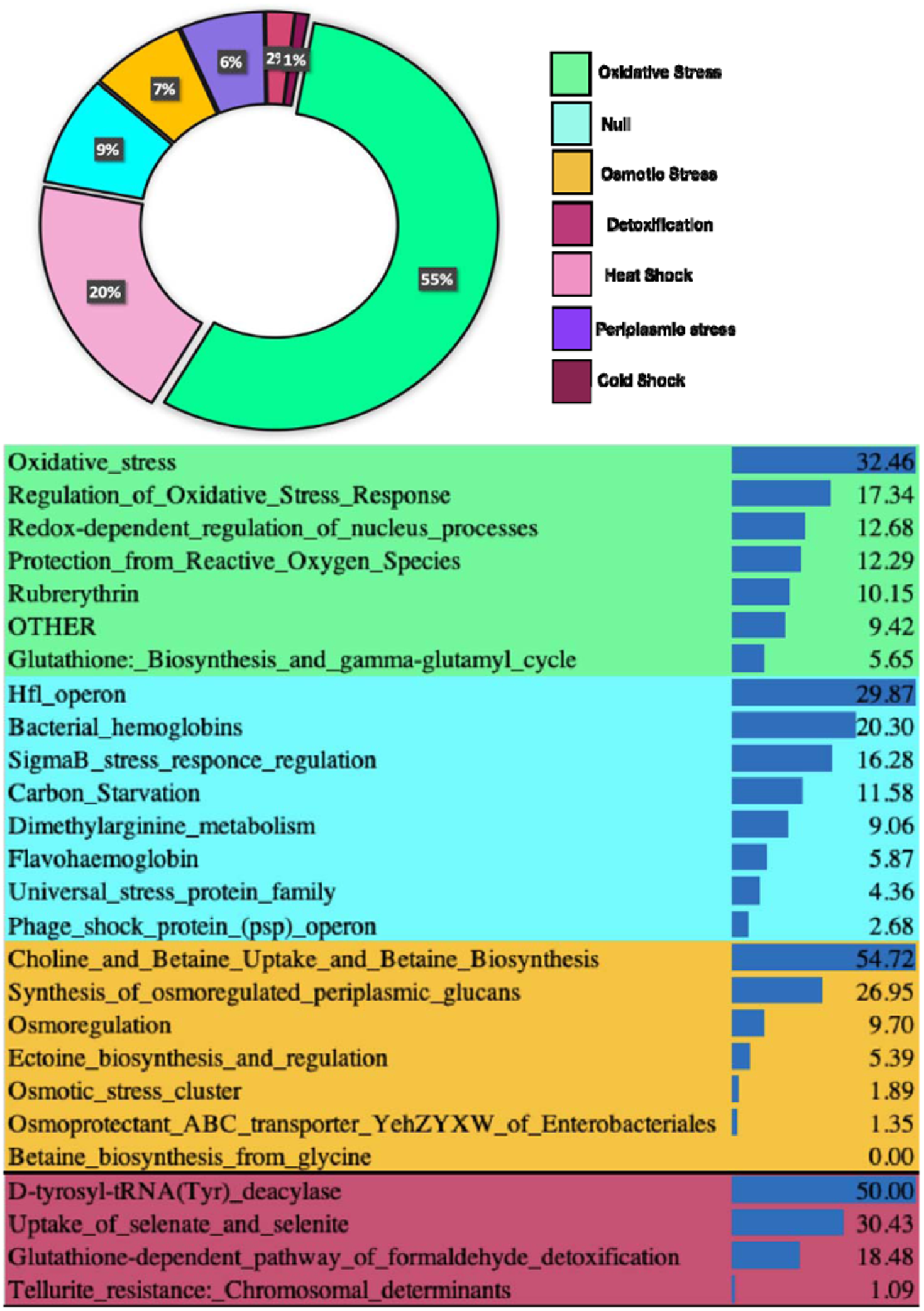
Relative differential abundance of functional features for various pathways for Stress related genes (Level-2 and Level-3) annotated from metagenome of polluted soils of JIE. The functions were mapped on Subsystem database on MG-RAST server with default parameters.

Most of these genes were found to be annotated from species of *Vibrio, Bacteroides, Acinetobacter, Pseudomonas, Desulfovibrio, Clostridium, Shewanella, Pelobacter, Bacillus, Desulfomicrobium, Desulfococcus, Thermoanaerobacter, Geobacter, Thermotoga, Marinobacter, Parabacteroides, Escherichia, Rhodobacter, Burkholderia, Petrotoga, Parvibaculum, Nitrosococcus, Desulfuromonas, Sulfurimonas, Candidatus, Arcobacter* etc. Various stress response enzymes like Catalases, Peroxidases, glutathione-S-transferases were annotated for polluted site of JIE.

#### 3.5.4 Metabolism of Carbon

The functionome from polluted metagenome of JIE was rich in diversity of genes for various carbon metabolizing pathways. The most abundant genes were annotated for central carbohydrate metabolism pathway when categorized at subsystem level-2. While delineating the scenario at subsystem level-3 revealed the most abundant functional features for Sugar utilization in Thermotogales, Serine-Glyoxylate cycle and TCA cycle with others.

The metabolic pathways for disaccharide, oligosaccharide and central metabolism of other carbon compounds are universally present in most of the microorganisms. However, few of the dominantly observed bacterial genera exhibiting various carbon metabolizing ability at polluted habitat of JIE were taxonomically assigned to *Geobacter* spp., *Desulfomicrobium* spp., *Thioflavicoccus* spp., *Thauera* spp., *Pseudomonas* spp., *Gluconacetobacter* spp., *Forficula* spp., *Sphaerochaeta* spp., *Clostridium* spp., *Petrotoga* spp., *Paracoccus* spp., *Clostrridium* spp., *Allochromatium* spp., *Mesotoga* spp., *Desulfosporosinus* spp., *Magnetospirillum* spp., *Anaeromyxobacter* spp., *Thioalkalivibrio* spp., *Hyphomicrobium* spp., *Haliangium* spp., *Desulfobacca* spp., *Streptosporangineae* spp., *Azospirillum* spp., *Rhizobium* spp., *Rhodobacter* spp., *Desulfobulbus* spp., *Arcobacter* spp., *Azoarcus* spp., *Thiocystis* spp., *Symbiobacterium* spp., *Dinoroseobacter* spp., *Desulfarculus* spp., *Bacillus* spp., *Parvibaculum* spp., *Halomonas* spp., *Desulfococcus* spp., *Sphingobium* spp. and many more.

#### 3.5.5 Antibiotic Resistance and Heavy Metal Resistance

The most abundant category under virulence disease and defence was annotated for antibiotic resistance and heavy metal resistance category. Various genes related to Cobalt-Zinc-Cadmium resistance, multidrug resistance efflux, resistance to fluoroquinolones. Genes related to Copper homeostasis, mercury resistance, Arsenic resistance, resistance to chromium resistance were annotated (Devpura et al., 2017). The taxonomic players for this were identified mainly from species of *Bacteroides, Vibrio, Pseudomonas, Acinetobacter, Desulfovibrio, Shewanella, Clostridium, Pelobacter, Geobacter, Escherichia, Marinobacter, Parabacteroides, Thiobacillus, Desulfococcus, Syntrophus, Parvibaculum, Desulfomicrobium, Sulfurimonas, Sulfurovum, Alkaliphilus, Burkholderia, Anaeromyxobacter, Syntrophobacter, Arcobacter* and *Dinoroseobacter.* The genes were categorised as subsystem level-2 and further studied at subsystem level-3 which can be understood from Figure 16.

**Figure 16:**
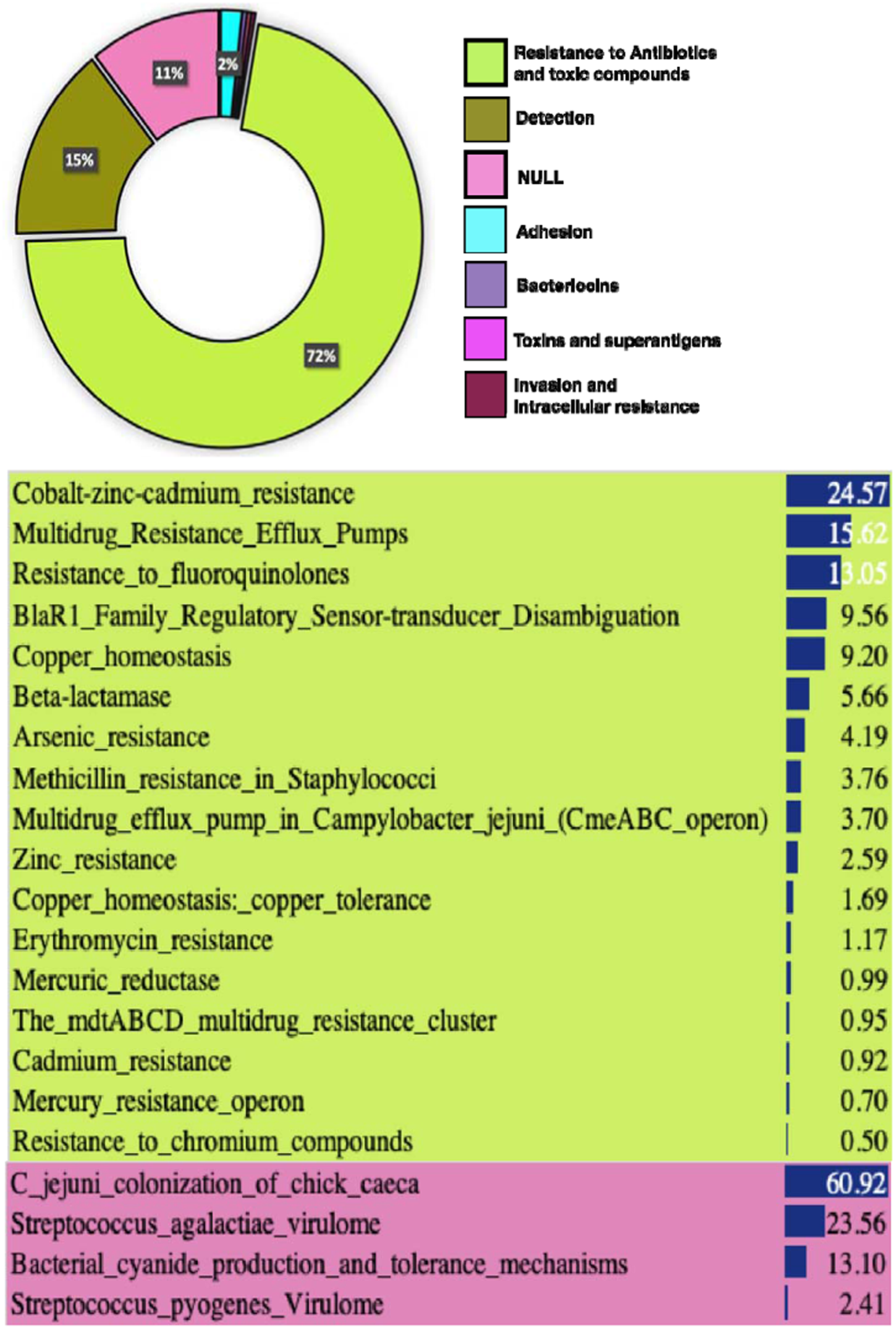
Relative differential abundance of functional features for various pathways for resistance towards heavy metals, antibiotics and drugs (Level-2 and Level-3), annotated from metagenome of polluted soils of JIE. The functions were mapped on Subsystem database on MG-RAST server with default parameters.

#### 3.5.6 Degradation of Aromatic Compounds

The catabolism of aromatic compounds by prokaryotes can be broadly classified into three categories: (i) peripheral (upper) pathways for the catabolism of aromatic compounds, (ii) metabolism of central (lower) aromatic intermediates and (iii) anaerobic degradation of aromatic compounds. The microbial catabolism (degradation) of aromatic compounds is a multistep process. This interrelated process involved dealkylation, deamination, decarboxylation, dechlorination, dehydrogenation, dioxygenation, hydrogenation, hydrolysis, hydroxylation, ortho-ring cleavage reactions, meta-ring cleavage reactions, phenyl ring cleavage reactions, s-triazine ring cleavage reactions and many others. These and other such reactions involved in xenobiotic catabolism (of catechol, biphenyl, naphthalene, phthalate, fluorobenzoate, xylene, phenylpropanoate, phenol and benzene amongst the other substituted and non-substituted aromatic compounds) are catalysesd by 1,2/2,3-dioxygease, decarboxylase, dechlorinase, dehalogenase, dehydrochlorinase, dehydrogenase, dioxygenase, hydrolase, reductase, monoxygenase, ligase, decarboxylating dehydrogrnases and many more.

These compounds are widely used for the production of dyes and dyes intermediates, pesticides, lubricants, rubbers, etc. The detail analysis of biodegradation genes (BDGs) displayed a marked competence of autochthonous microbial community of polluted ecosystem of JIE in catabolism of aromatic and other xenobiotic SOCs. In peripheral (upper) pathways, genes encoding the enzymes and proteins for catabolism of benzoate, biphenyl, chlorobenzoate and other chloroaromatic compounds, naphthalene and antracene, n-Phenylalkanoic acid, phenol, phenylpropanoid compounds, p-Hydroxybenzoate, Quinate, Salicylate ester toluene, etc. were annotated. In benzoate catabolism, genes for benzoate transport protein, benzoate MFS transporter (*benK*), Muconolactone isomerase [EC: 5.3.3.4] and Benzoate 1,2-dioxygenase [EC: 1.14.12.10] and Benzoylformate decarboxylase [EC: 4.1.1.7] were annotated. For chlorobenzoate and other chloroaromatic compounds catabolism, 3-oxoadipate CoA-transferase [EC: 2.8.3.6], Betaketoadipyl CoA thiolase [EC: 2.3.1.-], 2-chlorobenzoate 1,2-dioxygenase [EC: 1.14.12.13] were annotated. For phenol and phenylpropanoid compounds catabolism, phenol hydroxylase, Phenylpropionate dioxygenase and related ring-hydroxylating dioxygenases, Vanillate Odemethylase oxidoreductase [EC: 1.14.13.-], etc. were annotated.

In central (lower) pathways for aromatic intermediates, genes encoding for enzymes and proteins for catabolism of 4-Hydroxyphenylacetic acid, Catechol degradation via β-ketoadipate and meta/orthto-cleavage pathways, homogentisate pathways for aromatic compounds, Nheterocyclic aromatic compounds, Protocatechuate degradation via β–ketoadipate pathway, Salicylate and gentisate catabolism and other aromatic compounds were annotated. For 4-Hydroxyphenylacetic acid catabolism, 5-carboxymethyl-2-hydroxymuconate delta-isomerase [EC: 5.3.3.10], 2-hydroxyhepta-2,4-diene-1,7-dioate isomerase [EC 5.3.3.-] were annotated. For catabolism of aromatic compounds through meta-cleavage pathway, 2-hydroxy-6-oxo-6- phenylhexa-2,4-dienoate hydrolase [EC: 3.7.1.-], 2-polyprenylphenol hydroxylase and related flavodoxin oxidoreductases, Acetaldehyde dehydrogenase/acetylating [EC 1.2.1.10] (also involved in gene cluster for degradation of phenols, cresols, catechol), 2-hydroxymuconic semialdehyde hydrolase [EC: 3.7.1.9], Catechol 2,3-dioxygenase [EC: 1.13.11.2], 3,4- dihydroxyphenylacetate 2,3-dioxygenase, Protocatechuate 4,5-dioxygenase [EC: 1.13.11.8] were annotated. In catechol degradation through meta-cleavage pathway (catechol to acetyl- CoA / 4-methylcatechol to propanoyl-CoA) catechol 2,3-dioxygenase (*catE*) [EC:1.13.11.2], 4-hydroxy 2-oxovalerate aldolase (*mhpE*) [EC:4.1.3.39], and through ortho-clevage pathway (catechol to 3-oxoadipate), 3-oxoadipate enol-lactonase (*pcaD*) [EC:3.1.1.24], muconate cycloisomerase (*catB*) [EC:5.5.1.1] were annotated.

In Homogentisate pathway for aromatic compound degradation, 4-hydroxyphenylpyruvate dioxygenase [EC: 1.13.11.27], Aromatic-amino-acid aminotransferase [EC: 2.6.1.57] were annotated. For Protocatechuate degradation via β-ketoadipate_pathway, β-ketoadipate enollactone hydrolase [EC: 3.1.1.24],4-carboxymuconolactone decarboxylase [EC: 4.1.1.44], Protocatechuate 3,4-dioxygenase [EC: 1.13.11.3], 3-oxoadipate enol-lactone hydrolase, etc. were annotated. In salicylate and gentisate catabolism, Maleylacetoacetate isomerase [EC: 5.2.1.2], Fumarylacetoacetase [EC: 3.7.1.2], Gentisate 1,2-dioxygenase [EC: 1.13.11.4] were annotated.

For catabolism of aromatic compounds under anaerobic and/or micro-aerophilic environment, genes encoding for enzymes and proteins for benzoate, toluene, ethylbenzene catabolism were annotated. In benzoate catabolism under anaerobic conditions, Benzoyl-CoA reductase (*badDEG*) [EC: 1.3.99.15], Benzoate-CoA ligase [EC: 6.2.1.25], Glutaryl-CoA dehydrogenase [EC: 1.3.99.7] were annotated. For toluene and ethylbenzene degradation, Acetophenone carboxylase, Ethylbenzene dehydrogenase [EC: 1.17.99.2], Succinyl-CoA:(R)-benzylsuccinate CoA-transferase [EC: 2.8.3.15] were annotated.

Genes encoding for 4-hydroxyphenylacetate 3-monooxygenase [EC: 1.14.13.3], Nitrilotriacetate monooxygenase [EC: 1.14.13.-] (involved in aromatic amin catabolism), Shikimate kinase I & II [EC: 2.7.1.71], 4-oxalocrotonate decarboxylase [EC: 4.1.1.77], 4-hydroxybenzoyl-CoA reductase [EC: 1.3.99.20], (involved in benzoate degradation), 4-cresol dehydrogenase (hydroxylating) flavoprotein [EC: 1.17.99.1] (cresol degradation), Fumarylacetoacetate hydrolase family protein (Gentisare degradation) were also annotated. Similarly, genes for benzoate/toluate 1,2-dioxygenase (*benA-xylX*) [EC:1.14.12; 1.14.12.-], dihydroxycyclohexadiene carboxylate dehydrogenase (benD-xylL) [EC:1.3.1.25; 1.3.1.-] involved in benzoate, fluorobenzaote, xylene and other substituted single ring aromatic compounds were annotated. Protocatechuate-3,4-dioxygenase (*pcaGH*) involved in benzoate, polycyclic aromatic compound catabolism were also annotated.

In aminobenzoate catabolism, 4-nitrophenyl phosphatase [EC: 3.1.3.41], amidase [EC: 3.5.1.4], (hydroxyamino) benzene mutase (*hab*) [EC: 5.4.4.1], 4-hydroxybenzoate decarboxylase (*bsdC*) [EC: 4.1.1.61] were annotated. For Nitrotoluene and chloroalkane/chloroalkene degradation, nitroreductase (*nfsA*) [EC: 1.-.-.-], dihydropteridine reductase (*nfnB*) [EC: 1.-.-.-, 1.5.1.34] and haloalkane dehalogenase (*dhaA*) [EC: 3.8.1.5] were annotated respectively. In aminobenzoate catabolism, vanillate monooxygenase (*vanA*) [EC: 1.14.13.82], 4-hydroxybenzoate decarboxylase (*bsdC*) [EC: 4.1.1.61] and for Styrene degradation, homogentisate-1,2-dioxygenase (*hgd*) [EC: 1.13.11.5], glutaconate CoAtransferase, (*gctA*) [EC: 2.8.3.12] were annotated.

Functions associated with catabolism of aromatic compounds were also annotated from pristine habitat (although with variable gene count). Since enzymes involved in catabolism of xenobiotic aromatic compounds are highly non-specific, with their actual biological roles in microbial physiology/metabolism are different, they can also catabolise synthetic organic carbons when present in the environment. Moreover, at pristine ecosystem, natural compounds like lignin are degraded by such enzymes. A broad description of this can be seen in Figure 17. Genes encoding the enzymes like, benzoate-CoA ligase, benzoyl-CoA reductase and benzoyl-CoA dihydrodiol lyase, catalyses the degradation of aromatic compounds under anaerobic conditions and significantly annotated from polluted habitat.

**Figure 17:**
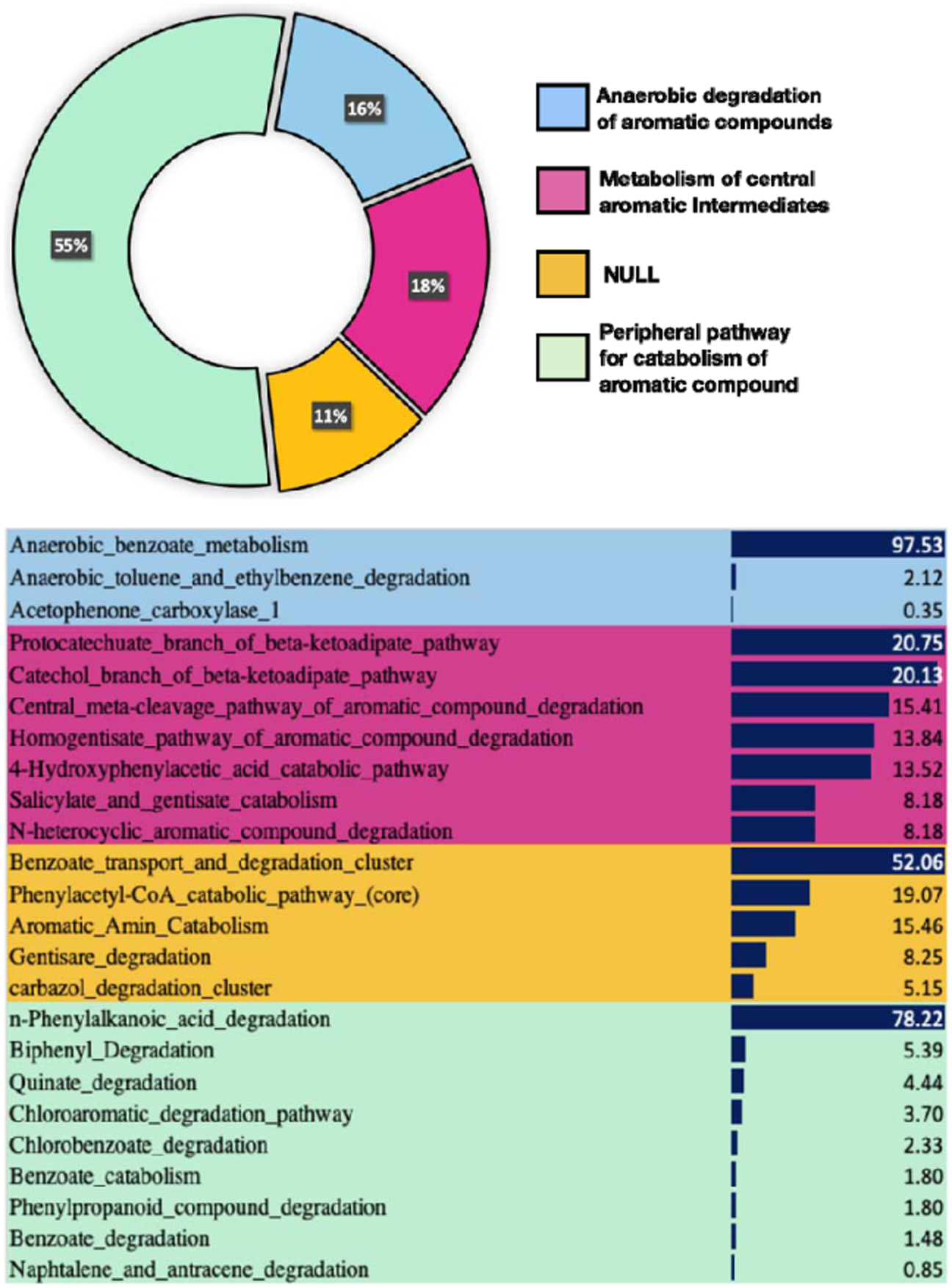
Relative differential abundance of functional features for various pathways for resistance towards heavy metals, antibiotics and drugs (Level-2 and Level-3), annotated from metagenome of polluted soils of JIE. The functions were mapped on Subsystem database on MG-RAST server with default parameters.

During anaerobic catabolism of benzoate, benzoate-CoA ligase catalyses the conversion of benzoate into benzoyl-CoA, which inturn converted into cyclic-1,5-dienoyl-CoA by ATP independent pathway (Cabral and Martins, 2018). The dearomatization of benzoyl-CoA under anaerobic metabolism was catalyzed by benzoyl-CoA reducatse (Fuchs et al., 2011). Few facultative anaerobes, also initiates the aerobic catabolism of benzoate through benzoate-CoA ligase (Carmona et al., 2009).

## 4.0 Discussion

In one of the microbial community studies on wastewater treatment plant, Muszynski et al. (2015) found the dominance of Actinobacteria, β-proteobacteria and Chloroflexi across the three seasons. In a similar study at wastewater treatment plant with anoxic-aerobic membrane bioreactor (MBR), Phan et al. (2016) observed that Methylophilales, Myxococcales, Rhodocyclales and phylum Plantomycetes were playing active role in removing trace organic matters along with total organic carbon and nitrogen. In one of our earlier studies on polluted aquatic environment of Alang-Sosiya ship breaking yard, near Bhavnagar, the ship breaking activities has released large amount of petroleum hydrocarbons, polycyclic aromatic hydrocarbons, heavy metals and other xenobiotic compounds. Because of this chronic release, the bacterial community shifted from a dominance of Gammaproteobacteria (at pristine sites) towards Epsilonproteobacteria at polluted sites, where Proteobacteria, Firmicutes, and Bacteroidetes were predominantly found (Patel et al., 2014). In this study also a marked changed in bacterial diversity between polluted and pristine sites has been found.

The active microbial assemblage of JIE shows the abundance of anaerobes, thermophile and sulfate reducers as well as methanogens like *Petrotoga, Desulfovibrio, Thermotoga, Methanoculleus, Methanosaeta* etc. These microbes have been extensively reported for their niche specific roles in various co-metabolic degradation processes in studies performed to understand oil, diesel and other xenobiotic biodegradation potential. (Lien et al., 1998; Yamane et al., 2008; Davey et al., 2015; Dorosh et al., 2016; Pradhan et al., 2016; Singh et al., 2018; Zhou et al., 2017; Wolters et al., 2018; Gao et al., 2019; Liao et al., 2019; Chen et al., 2019). The archaeal diversity from our study has found to play important role in biostimulation, bioaugmentation and bioremediation studies from contaminated environments (Roy et al., 2018; Krzmarzick et al., 2018; Bharadwaj, 2018).

Industrial pollution may adversely affect the autochthonous microbiome of terrestrial and aquatic ecosystem equally. The earlier studies initiated in pure culture era and later on by 16S rRNA gene cloning approach did provide cell number estimation and along with microscopic methods (e.g. FISH) and supported perception of community shift induced due to regional pollution. Nevertheless, with the technology advancement and in particular NGS platforms has completely changed our perception of microbial diversity dynamics. However, mathematical and statistical calculations provided us depth and breadth of microbial community of terrestrial and aquatic ecosystem; it was after the advancement in sequencing technologies, the true nature of microbial community was actually realized. Many microbial ecological models have been revisited, in NGS era. But on contrary, it has been now revealed that polluted environment is biologically very active than some of the non-polluted/pristine habitats and hosts finely tuned and well defined sets of microbes (Bargiela et al., 2015; Devpura et al., 2017). The ecological phylotypic richness of such habitats depends on chemical nature of synthetic organic compounds (SOCs) and other xenobiotic pollutants present in the ecosystem. Moreover, it is not only the nature of SOCs, other environmental (abiotic and biotic) factors have shown to define the polluted microbiome community (Kleinsteuber et al., 2012). Nevertheless, it is a matter of further study how multiple environmental factors (which make the polluted ecosystem highly heterogeneous) affect and impact the autochthonous community and which of individual or sets of factors are driving force determining the final catabolic phenotype of polluted environment (Kleinsteuber et al., 2012).

In this study, we could assign many of taxa to the micro-niches on the basis of results obtain by amplicon profiling, functional co-relation (re-construction) of metabolic pathways and predicted metaphenomic study. The exercise allows us to infer and describe the theoretical ecological model of microbial (bacterial) ecosystem of polluted environment of polluted river flowing across the Jetpur Industrial Estate. The infer microbial ecological model coupling with Carbon, Nitrogen, Sulphur biotransformation and catabolism of infused SOCs, allow us to segregate the phenotype into obligate/facultative aerobic, anoxic, anaerobic niches. Through this model an extensive metabolic pathway prevailing at polluted ecosystem was elucidated. But, since the proposed ecological model is based on amplicon and metagenomic content and predictive metaphenomics and their differential abundance of the genome (in a community context), it requires further validation through e.g. metagenomics, metaproteomics and other relevant experimental methods (Zhang et al., 2019; Schiebenhoefer et al., 2019).

There are several geographical regions across the globe, perturbed with anthropogenic activities. Amongst many of them, is industrially polluted terrestrial and aquatic ecosystems affected by industrial activities, which remains poorly understood on a large scale. Here, firstly we have compared the influence of geographical and environmental constrains along with anthropogenic forces (i.e. infusion of SOCs, heavy metals and other xenobiotic compounds) on bacterial communities, at polluted as well as non-polluted areas of JIE. The results pointed to few signatures functional characteristics of polluted microbiome. Since we could not measure the bacterial diversity before the on-set of pollution, the bacterial diversity was compared with pristine conditions prevailing at the time of sampling near the polluted sites. The polluted ecosystems like JIE comprises of water logging zones (similar to intertidal zones of marine coastal ecosystem), which promotes regular fluctuations in environmental conditions including oxygen and nutrient availability. The biotic and abiotic ecological interactions together with physico-chemical heterogeneity of the perturbed ecosystem, contributes to catabolic efficiency of autochthonous microbial community, which needs to alternate between aerobic and anaerobic mode of respiration and catabolism of recalcitrant SOCs (Cabral and Martins, 2018; Dixit et al., 2015; Kleinsteuber et al., 2012). Genes encoding the enzymes (and proteins) involved in catabolism of aromatic compounds under aerobic conditions i.e. aerobic pathways of aromatic compound degradation (meta- and ortho-pathways/cleavage), like monooxygenase and dioxygenase, (catechol-2,3-dioxygenase), 3-oxoadipate CoA-transferase, β-ketoketoadipase enol-lactone hydrolase were annotated in higher significant number suggests many of the aromatic compounds were being catabolise under aerobic conditions at polluted habitat of JIE by autochthonous microbial community.

Under the influence of microbial metabolism, aromatic compounds are first transformed from relatively inter into more reactive intermediary compounds such as catechol, protocatechol and their derivatives, followed by cleavage of intradiol or extradiol rings by molecular oxygen (i.e. ring opening reactions) (D’Argenio et al., 2014).

Catechol and its derivatives are the common central intermediary compounds in the aromatic catabolic/metabolic pathways leading to complete mineralization via ortho- or central or meta-cleavage pathways (Cabral and Martins, 2018; Nair et al., 2008). The prokaryotic metabolism of aromatic compounds through aerobic (upper) pathways leads to catechol degradation pathways. One of the significant enzymes in the reaction is catechol-2,3-dioxygenase, catalyzes aerobic degradation/catabolism of several aromatic compounds such as benzoate, salicylate, phenol and other single ring compounds. Another enzyme catechol-1,2-dioxygenase catalyses ring opening reactions and convert aromatic compounds into cis, cis-mucoante. The polluted habitats of JIE are evidently and preferentially anaerobic and same has been postulated and predicted during metaphenomic study. However, the root activities of plants, movement of stray domestic animals and routine input of liquid wastes bring oxygen to the micro-niches at polluted ecosystem. The underlying soil and sediments are waterlogged, but are periodically exposed to the open environment (Xin et al., 2013). Moreover, the upper top surface soils due to regular water movement and other activities become porous and facilitates water percolation and sparse aeration (Hossain and Nuruddin, 2016).

Gene encoding the important enzymes/proteins, 4-hydroxybenzoate transporter and benzoate transporter proteins were significantly annotated from polluted habitats. This enzyme (4-hydroxybenzoate transporter) is a permease (*pcaK*), which catalyzes the transport of aromatic acids like 2,4-dihydroxybenzoate, salicylate, 3-hydroxybenzaote, protocatechuate, gentisate, vanillate into the cell (Pernstich et al., 2014; Nichols and Harwood, 1997). 4-hydroxybenzoatetransporter is encoded as a part of aromatic acids catabolising during β-ketoadipate pathway and has two physiological functions: chemotaxis and transport (Luu et al., 2015; Nichols and Harwood, 1997). The β-ketoadipate pathway is thought to be central most pathway form aromatic degradation and is widely disturbed into soil bacterial and fungi (Fuchs et al., 2011). It is involved in catabolism of many toxic aromatic SOCs and transformed into non-toxic simpler compounds which enters in central TCA cycle and other anabolic metabolism (Wells and Ragauskas, 2012).

Genes encoding the enzymes like, benzoate-CoA ligase, benzoyl-CoA reductase and benzoyl-CoA dihydrodiol lyase, catalyses the degradation of aromatic compounds under anaerobic conditions and significantly annotated from polluted habitat. During anaerobic catabolism of benzoate, benzoate-CoA ligase catalyses the conversion of benzoate into benzoyl-CoA, which in turn converted into cyclic-1,5-dienoyl-CoA by ATP independent pathway (Cabral and Martins, 2018). The dearomatization of benzoyl-CoA under anaerobic metabolism was catalyzed by benzoyl-CoA reductase (Fuchs et al., 2011). Few facultative anaerobes, also initiates the aerobic catabolism of benzoate through benzoate-CoA ligase (Carmona et al., 2009).

## 4.0 Conclusion

The study to identify the microbial lifestyle of sediments from Bhadar River flowing across Jetpur Industrial Estate revealed a substantial intelligence of microbes in response to press disturbances. The substantial increase in anaerobic niche specific metabolic capabilities because of chronic infusion of various dyes and other synthetic organic compounds shows the remarkable metabolic plasticity of autochthonous microbiome at polluted sites of JIE. A prevalence of Sulfate reducing bacteria (SRBs), methane metabolizing microbes and fermentative microbes shows that the various organic chemicals were metabolized and transformed by syntrophic interactions among all these microbes. This lifestyle change, shifting from aerobic respiration towards sulfur respiratory mechanisms, appears to be a highly niche-specific phenomenon. In our study we can conclude that habitat selection drives the microbial assembly and their lifestyle at polluted site from Jetpur Industrial Estate.

## Acknowledgements

Authors are grateful Department of Biotechnology (DBT), and Department of Science and Technology (DST), Ministry of Science and Technology, New Delhi, India. ND duly acknowledges University Grants Commission, New Delhi, India for financial support in form of Junior Research Fellowship.

## Authors Contribution

ND: Investigation, Methodology, Writing – original draft, Visualization, Data curation. KRJ: Methodology, Investigation, Validation, Visualization, Writing – original draft, Writing – review & editing, Data curation, Formal analysis, Conceptualization, Project administration. DM: Writing – review & editing, Validation, Supervision, Resources, Project administration, Funding acquisition, Conceptualization.

